# Inhibition of the BMP pathway suppresses tumor growth via downregulation of EGFR in MEK/ERK-dependent colorectal cancer

**DOI:** 10.1101/2022.11.01.514643

**Authors:** Shota Shimizu, Jumpei Kondo, Kunishige Onuma, Kasumi Ota, Mayumi Kamada, Yohei Harada, Yoshihisa Tanaka, Mai Adachi Nakazawa, Yoshinori Tamada, Yasushi Okuno, Kenji Kawada, Kazutaka Obama, Robert J Coffey, Yoshiyuki Fujiwara, Masahiro Inoue

**Author notes:** **Correspondence** Jumpei Kondo, Department of Clinical Bio-resource Research and Development, Graduate School of Medicine Kyoto University, 46-29 Shimoadachi-cho, Sakyou-ku, Kyoto, 606-8501, Japan., Department of Molecular Biochemistry and Clinical Investigation, Division of Health Science, Osaka University Graduate School of Medicine, 1-7 Yamadaoka, Suita City, Osaka, 565-0871, Japan.

## Abstract

The bone morphogenetic protein (BMP) pathway promotes differentiation and induces apoptosis in normal colorectal epithelial cells. However, the effect of the BMP pathway in colorectal cancer (CRC) is controversial; it can either be tumor promoting or tumor suppressing, depending on the study. In this study, we found that CRC cells reside in a BMP-rich environment based on RNA-sequencing database analysis. Suppression of BMP using a specific BMP inhibitor, LDN193189, suppresses the growth of organoids in some CRC cases. CRC organoids treated with LDN193189 exhibited a decrease in epidermal growth factor receptor, which was, at least in part, mediated by protein degradation induced by leucine-rich repeats and immunoglobulin-like domains protein 1 (LRIG1). Among CRC organoid panels from 18 different patients, suppression of organoid growth by BMP inhibition correlated with the induction of *LRIG1* gene expression. Notably, knockdown of LRIG1 in organoids diminished the growth-suppressive effect of LDN193189. Furthermore, simultaneous treatment with LDN192189 and trametinib, an FDA-approved MEK inhibitor, resulted in a combination effect in both *in vivo* and *in vitro* xenograft tumor treatment in CRC organoids, which are susceptible to growth suppression by LDN193189. Taken together, the simultaneous inhibition of BMP and MEK can be a novel treatment option in CRC cases, and evaluating *in vitro* growth suppression and *LRIG1* induction by BMP inhibition using patient-derived organoids could offer functional biomarkers for predicting potential responders.

## 1. Introduction

Bone morphogenetic protein (BMP) ligands are members of the transforming growth factor beta (TGF-β) superfamily, which trigger intracellular signaling via specific receptors upstream of common SMAD4 proteins. TGF-β has been extensively studied for its role in cancer pathogenesis, including colorectal cancer (CRC), especially in the epithelial–mesenchymal transition (EMT) ^1,2^. However, the role of BMP ligands in cancer has been studied less extensively. In the normal colorectal epithelium, BMPs 2, 4, and 7 are secreted from stromal cells, such as myofibroblasts, which promote the differentiation of epithelial cells and suppress the stemness of epithelial stem cells at the crypt base ^3,4^. In CRC, BMPs are reported both as tumor suppressing and promoting molecules, depending on the study. Beck et al. reported that BMP signaling is intact and suppresses the growth of human colon cancer cell lines ^5^. Lombardo et al. revealed that BMP4 induces differentiation of human CRC stem cells ^6^. In contrast, Lorente-Trigos et al. reported that BMP signaling promotes the growth of xenograft tumors derived from primary human CRC ^7^. More recently, Yokoyama et al. reported that autocrine BMP4 signaling is a therapeutic target in CRC cell lines ^8^. In addition to the context dependencies of the BMP effect that have been discussed in various cancers ^9,10^, these controversies might represent inter-tumor heterogeneity among the CRC population. One possible explanation for the latter is that the mutation status of *SMAD4* and *TP53* is suggested to determine cell fate by BMP ^11^. However, further studies on the heterogeneous effects of BMPs are warranted.

Epidermal growth factor receptor (EGFR) is a receptor tyrosine kinase important for the survival and proliferation of intestinal and colorectal epithelial cells. Therefore, the regulation of EGFR signaling is critical for the pathology of CRC. Indeed, molecular-targeted drugs for EGFR, such as cetuximab and panitumumab, are clinically used for treating advanced CRC with wild-type *KRAS* ^12^. We have reported that *KRAS* mutant CRC can be characterized into two groups: a partially responsive group, in which cetuximab had a substantial growth inhibitory effect, and a resistant group, in which no effect was observed ^13^. The CRC cells in the partially responsive group showed a combined effect of cetuximab and trametinib, a MEK inhibitor, both *in vitro* and *in vivo*. These observations support targeting EGFR, even in *KRAS* mutant CRC. However, selecting a sensitive case is essential because cancer is a highly heterogeneous disease.

To address the heterogeneity of cancer, more researchers have been utilizing the three-dimensional organoid culture technique in recent years. Patient-derived organoids retain the physiological features of parental cancer cells and reflect the heterogeneity of disease subgroups ^14^. We have reported and utilized a method for organoid preparation, the cancer tissue-originated spheroid (CTOS) method, which prepares cancer organoids with high purity, yield, and viability by retaining cell-cell interactions throughout the procedure ^14,15^. In this study, the effect of BMP inhibition was evaluated using a panel of patient-derived CRC organoids established using the CTOS method. A group of CRC organoids sensitive to BMP inhibition was further investigated for the mechanism of growth suppression, and the possibility of a combination therapy was explored.

## 2. Material and Methods

### 2.1 Patient samples and animal studies

This study was approved by the Institutional Ethics Committees at Osaka International Cancer Institute (1712225296, 1803125402) and Kyoto University (R1575, R2444) and was performed in accordance with the Declaration of Helsinki. The surgical specimens were obtained from Osaka International Center Institute and Kyoto University after obtaining written informed consent.

The animal studies were approved by the Institutional Animal Care and Use Committee of Kyoto University and Osaka International Cancer Institute and were performed according to institutional guidelines.

### 2.2 Generation and treatment of xenograft tumors

To expand the organoids, they were injected into NOD/SCID mice (CLEA Japan, Tokyo, Japan) to generate subcutaneous xenograft tumors. For the *in vivo* treatment studies, a mixture of 1,000 organoids in Matrigel (Corning Inc., Corning, NY, USA) was transplanted into the flank of BALB/cAJcl-nu/nu mice (CLEA Japan). LDN193189 (LDN, Sigma-Aldrich, St. Louis, MO, USA) solution was prepared in water and administered daily to mice at 3 mg/kg intraperitoneally. Trametinib solution was prepared in 0.5% methyl cellulose with 0.2% Tween-80 and administered using oral gavage at 0.3 μg/kg every other day. The subcutaneous xenograft volume was calculated using the formula: volume = (width)^2^ × (length)/2, and treatment was started when the volume exceeded 300 mm^3^. Mice with an excessive tumor volume (>2,000 mm^3^) were euthanized for ethical reasons.

### 2.3 Organoid preparation, culture, and cryopreservation

Organoid preparation of CRC patient tissue and xenograft tumors was performed using CTOS method as previously described ^13,15^. Briefly, the tumor specimens were mechanically and enzymatically digested, and the small fragments trapped by 100 μm or 40 μm cell strainers (BD Falcon, Franklin Lakes, NJ). Then, they were collected, cultured, and passaged in suspension in StemPro hESC (Invitrogen, Carlsbad, CA, USA) as three-dimensional organoids. Organoids were cryopreserved using CS10 (BioLife Solutions, Bothell, WA, USA), following the manufacturer’s instructions.

### 2.4 Vector construction and gene transfer

Tet-pLKO-puro was a gift from Dmitri Wiederschain (Addgene, plasmid #21915). Oligos for the leucine-rich repeats and immunoglobulin-like domains protein 1 *(LRIG1)* shRNA coding sequence (shLRIG1-Fw: CCGGTCCACACGGACCGCCTATAAACTCGAGTTTATAGGCGGTCCGTGTGGATTTTTG; shLRIG1-Rev: AATTCAAAAATCCACACGGACCGCCTATAAACTCGAGTTTATAGGCGGTCCGTGTGGA) were annealed and cloned into the tet-pLKO-puro vector using the AgeI and EcoRI cloning sites. shRNA constructs were introduced into organoids using lentiviral transduction with the spin infection technique ^16^. Organoids were selected with 2.0 μg/mL puromycin (InvivoGen, #ant-pr-1, San Diego, CA, USA), and the transcription of shRNAs was induced with 2.0 μg/mL doxycycline hyclate (Sigma-Aldrich, #D9891).

### 2.5 Organoid growth assay

Organoid growth was evaluated in growth factor–free (GF-free) medium (Advanced DMEM/F12 [Gibco, Thermo Fisher Scientific, Waltham, MA, USA] supplemented with 1x GlutaMax [Gibco]) or StemPro medium (StemPro hESC [Invitrogen]) containing 2.5% Matrigel GFR (BD Biosciences). Organoids were used for the growth assay 2 days after passage. Approximately 10 organoids (φ70–100 μm) per well in a 96-well plate were used for each condition. For the assays, LDN, trametinib (Selleck Chemicals, Houston, TX, USA), recombinant human EGF (PeproTech, Rocky Hill, NJ, USA), recombinant human heregulin (HRG, PeproTech), and bafilomycin A1 were added at the indicated doses and time points. To quantify the viability of organoids, an ATP assay was performed using CellTiter-Glo (Promega, Madison, WI, USA). Chemiluminescence values were obtained using a GloMax Discover Microplate Reader (Promega). Organoids were cultured for 5 days, and the relative ATP levels were adjusted with the initial numbers of organoids. GraphPad Prism 9 software (GraphPad Software, San Diego, CA, USA) was used to draw sigmoidal dose–response curves.

### 2.6 Statistical analysis

Significance was tested using an unpaired t-test for single comparisons and one-way ANOVA with post-hoc Tukey’s test for multiple comparisons. For correlation, Spearman’s rank correlation coefficient was designated as the r-value, and the significance of the correlation was evaluated using Spearman’s rank correlation test. Statistical significance was set at P < 0.05.

Additional information for materials and methods are described in Supporting Information (Doc S1).

## 3. Results

### 3.1 CRC cells reside in BMP ligand-rich environment

First, we profiled BMP pathway gene expression in CRC tissues using public databases. Using RNA-sequencing data from cancerous and normal tissues registered in the Cancer Genome Atlas (TCGA) ^17^ and normal tissues registered in Genotype-Tissue Expression (GTEx) ^18^, we found that in normal colorectal tissues, among the major *BMPs* (*2, 4, and 7*) of the intestine, *BMP4* and *BMP7* expressions were increased in CRC tissues. In contrast, *BMP2* expression was lower than that in normal tissues (Fig. 1A). In addition, single-cell RNA sequencing data were used to analyze the status of the BMP pathway by cell lineage, and analysis of pooled single-cell gene expression data from nine CRC patients ^19^ revealed that *BMP4* was mainly expressed in fibroblasts and epithelial tumor cells (Fig. 1B, 1C, S1). In contrast, *BMP7* was produced by various types of cells, including malignant cells and hematopoietic cells, such as T cells and B cells (Fig. 1B-C). The expression of ID1 and 2, major targets of the BMP/SMAD pathway, was upregulated in epithelial cell clusters, suggesting that the BMP pathway is active in cancer cells (Fig. 1D). These results suggest that the BMP pathway is involved in the development of the microenvironment in CRC, based on gene expression analysis of clinical tissue–derived samples.

**FIGURE 1.**
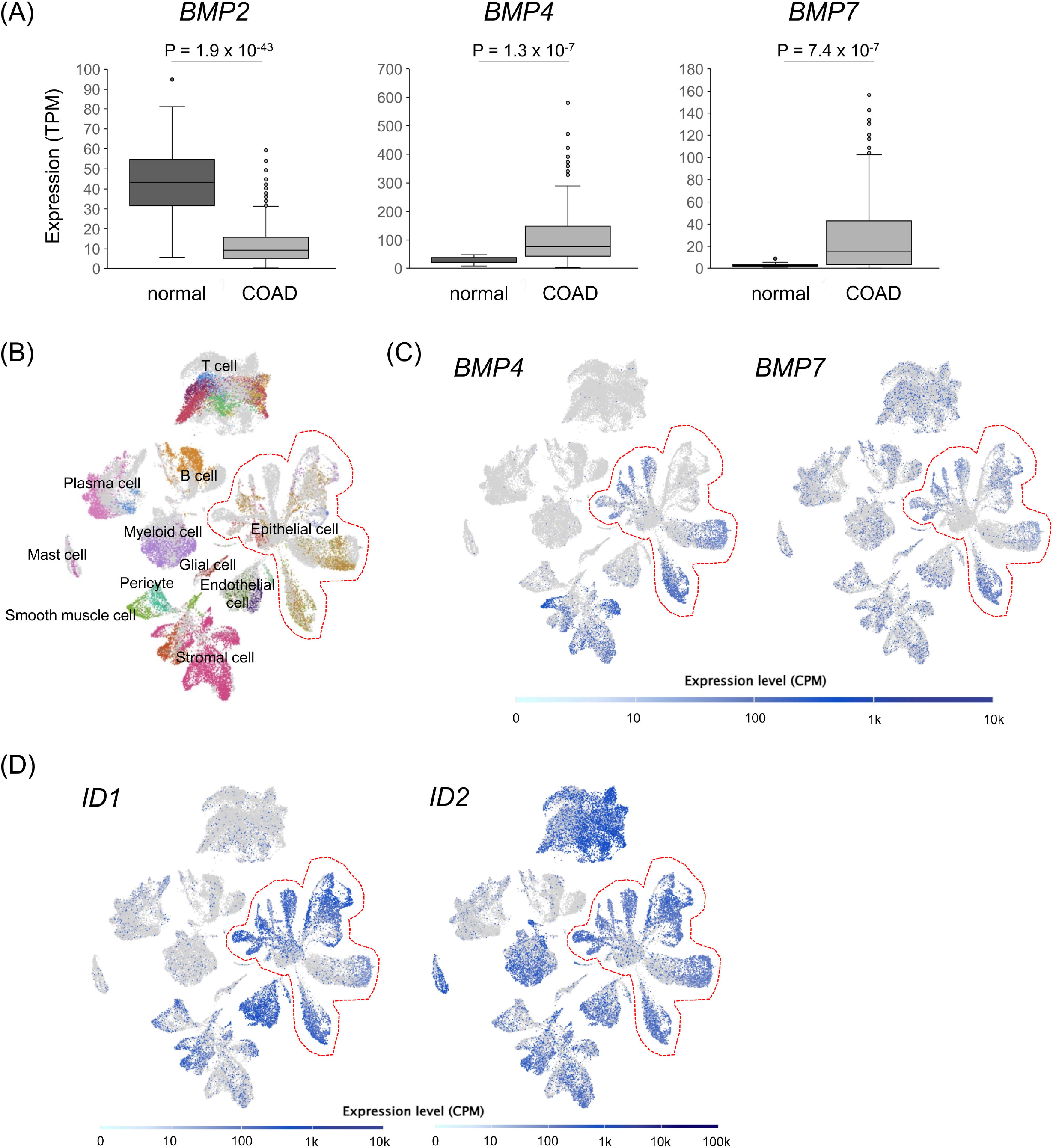
BMP pathway is active in clinical CRC tumors. **A**, Box-whisker plot of *BMPs 2, 4*, and *7* expressions (TPM) in normal colon (normal n = 41) and colorectal adenocarcinoma (COAD n = 308) tissue specimen. *P* values calculated by Student’s t-test are indicated. **B-D**, Uniform Manifold Approximation and Projection (UMAP) for the single-cell RNA sequencing data of 9 CRC patients’ specimens. Cell types corresponding to each cluster are designated on the UMAP (B). More detailed cluster labeling is presented in Fig. S1. *BMP4* and *BMP7* (C), as well as *ID1* and *ID2* (D) expression levels, are plotted on the UMAP. Areas enclosed by red dotted lines indicate the cluster of epithelial cells.

### 3.2 Inhibition of the BMP pathway suppresses CRC organoid growth with decreased EGFR

To investigate the role of the BMP pathway in CRC cells, the CRC organoid panel (Table S2) was tested for growth inhibition in a defined medium without growth factors using LDN, a BMP receptor inhibitor. We found that the effects of LDN varied among organoids from different cases (Fig. 2A-B). This phenomenon was confirmed with the recombinant BMP-inhibitory protein Noggin. The growth of C45 and CB3 LDN-sensitive organoids was significantly inhibited by Noggin, while that of C75 and C166 LDN-resistant organoids was not (Fig. S2). Next, the time course of the activation status of the signaling pathway was evaluated in these LDN-sensitive and LDN-resistant organoids. Phosphorylation of SMAD1/5, which are key signaling molecules of the BMP/SMAD pathways downstream of BMP receptors, increased over time in the absence of LDN (Fig. 2C), supporting the autocrine production of BMP ligands by CRC cells indicated from omics data (Fig. 1C). LDN suppressed SMAD1/5 phosphorylation, as expected (Fig. 2C). The level of EGFR, a major receptor tyrosine kinase in the pathogenesis of CRC ^20^, decreased in these CRC organoids treated with LDN, especially after long-term treatment (48 h, Fig. 2C). Phospho-ERK1/2 (pERK) was suppressed by LDN in C45, CB3, and C75 organoids but not in C166 organoids (Fig. 2C). Thus, suppression of EGFR/MEK/ERK by LDN was predicted to be the mechanism of growth inhibition. Therefore, we investigated the effect of MEK/ERK activation on LDN treatment. LDN treatment in the presence of a large excess of EGF (Fig. 2D) and HRG (Fig. 2E), a ligand of HER3, attenuated the growth suppression effect of LDN. Furthermore, the effect of LDN on the growth of CRC organoids was examined using a medium containing a cocktail of growth factors (IGF, HGF, and HRG) ^15,21^. Compared to the medium without growth factors (Fig. 2A), the growth inhibition by LDN was generally much lower (Fig. 2F). These results suggest that LDN inhibits CRC organoid proliferation via suppression of MEK/ERK through downregulation of EGFR as an upstream event.

**FIGURE 2.**
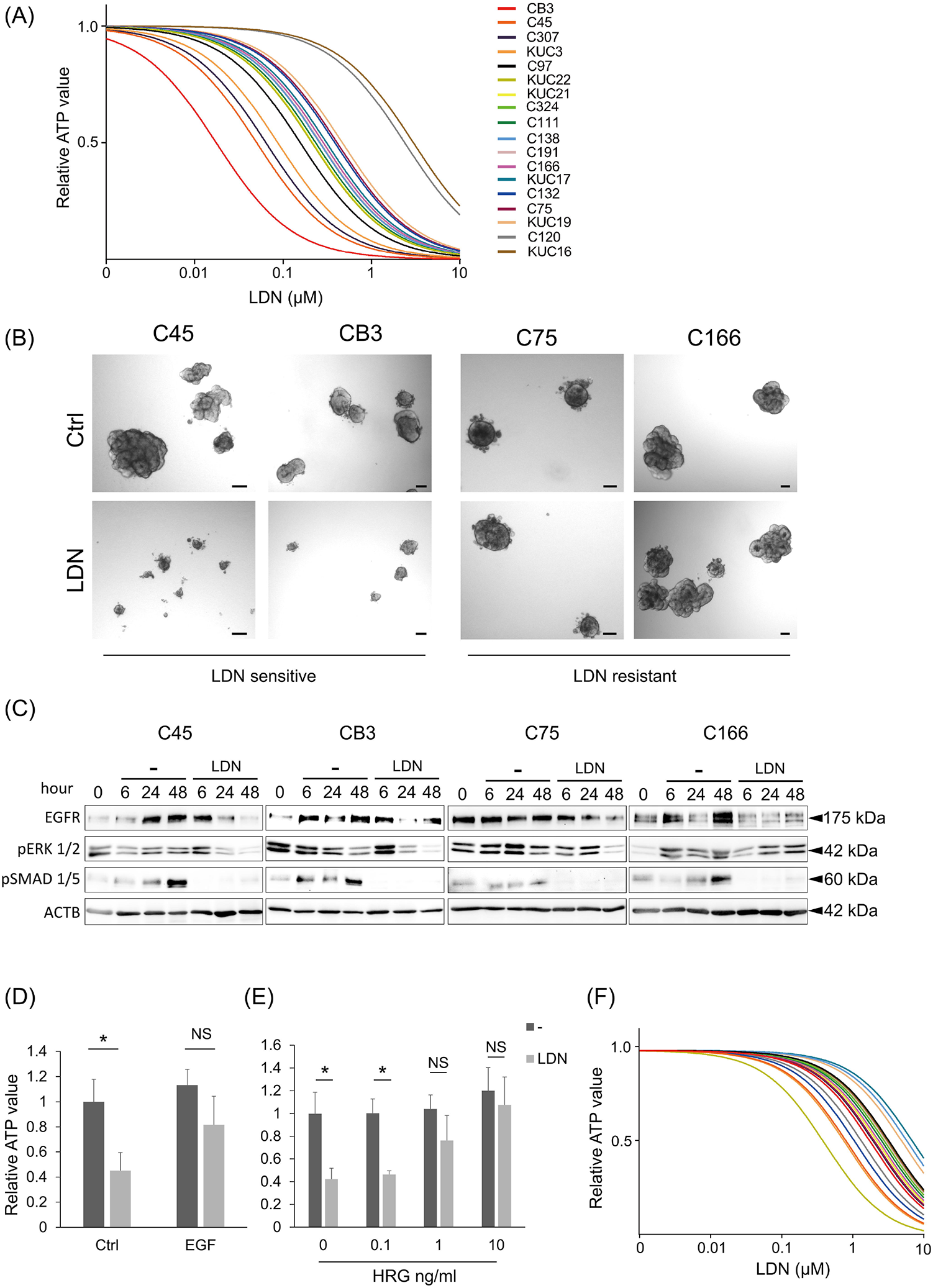
Inhibition of the BMP pathway suppresses CRC organoid growth. **A**, Dose-response curves of LDN in 18 CRC organoid lines cultured in the GF-free medium. **B**, Bright-field images of CRC organoids cultured in the presence (lower panels) or absence (Ctrl, upper panels) of 0.1 μM LDN for 5 days. Scale bars: 100 μm **C**, Immunoblotting analysis of CRC organoids cultured in the presence (LDN) or absence (-) of 0.1 μM LDN for the indicated time (hours). Proteins were detected with indicated antibodies. **D-E**, Cell viability assay results for C45 organoids. Organoids were treated with 0.1 μM LDN with or without 50 ng/mL EGF (N = 5 for each condition, D) or heregulin (HRG) at the indicated concentration (N = 3 for each condition, E). Data are presented as mean + SD. * P < 0.05, t-test with Bonferroni correction. **F**, Dose– response curves of LDN in 16 CRC organoid lines cultured in the StemPro hESC medium.

### 3.3 BMP inhibition induces LRIG1-mediated degradation of EGFR

Next, we investigated the mechanisms underlying the downregulation of the EGFR/MEK pathway by BMP inhibition. Despite the decreased EGFR protein levels, *EGFR* mRNA expression was only slightly downregulated in C45 and C75 organoids treated with LDN or Noggin and was not altered in C166 organoids (Fig. 3A). In addition, Noggin treatment upregulated the expression of *EGFR* in CB3 organoids (Fig. 3A). These results indicate that the suppression of organoid growth by BMP inhibition was not due to the transcriptional regulation of *EGFR*. We evaluated the induction of several other negative regulators of the EGFR/MEK pathway by BMP inhibition. EGFR is regulated at the post-translational level, and ligand-stimulated EGFR proteins are endocytosed either to be degraded or recycled ^22,23^. Some negative regulators of EGFR, including LRIG1 ^24^ and ERRFI1 (MIG-6) ^25^, are known to trigger this degradation machinery. *LRIG1* expression was highly upregulated in LDN-treated C45 and CB3 organoids and slightly upregulated in C75 organoids (1.47-fold) (Fig. 3B). However, *ERRFI1* was not upregulated in any of the organoids evaluated (Fig. S3). Lysosomal degradation, led by ubiquitination, has been reported as a mechanism by which LRIG1 degrades EGFR ^26,27^. Indeed, EGFR was highly ubiquitinated in the C45 organoid following LDN treatment (Fig. 3C). Moreover, bafilomycin A1, a lysosomal inhibitor, inhibited the downregulation of EGFR by LDN (Fig. 3D) and growth suppression by LDN (Fig. 3E). LRIG1, a pan-ErbB negative regulator, did not decrease the expression of HER2 and 3 proteins (Fig. 3F). These results indicate that EGFR degradation was involved in the LDN-induced growth suppression of CRC organoids and further suggest that LRIG1 induction is responsible for the LDN effect.

**FIGURE 3.**
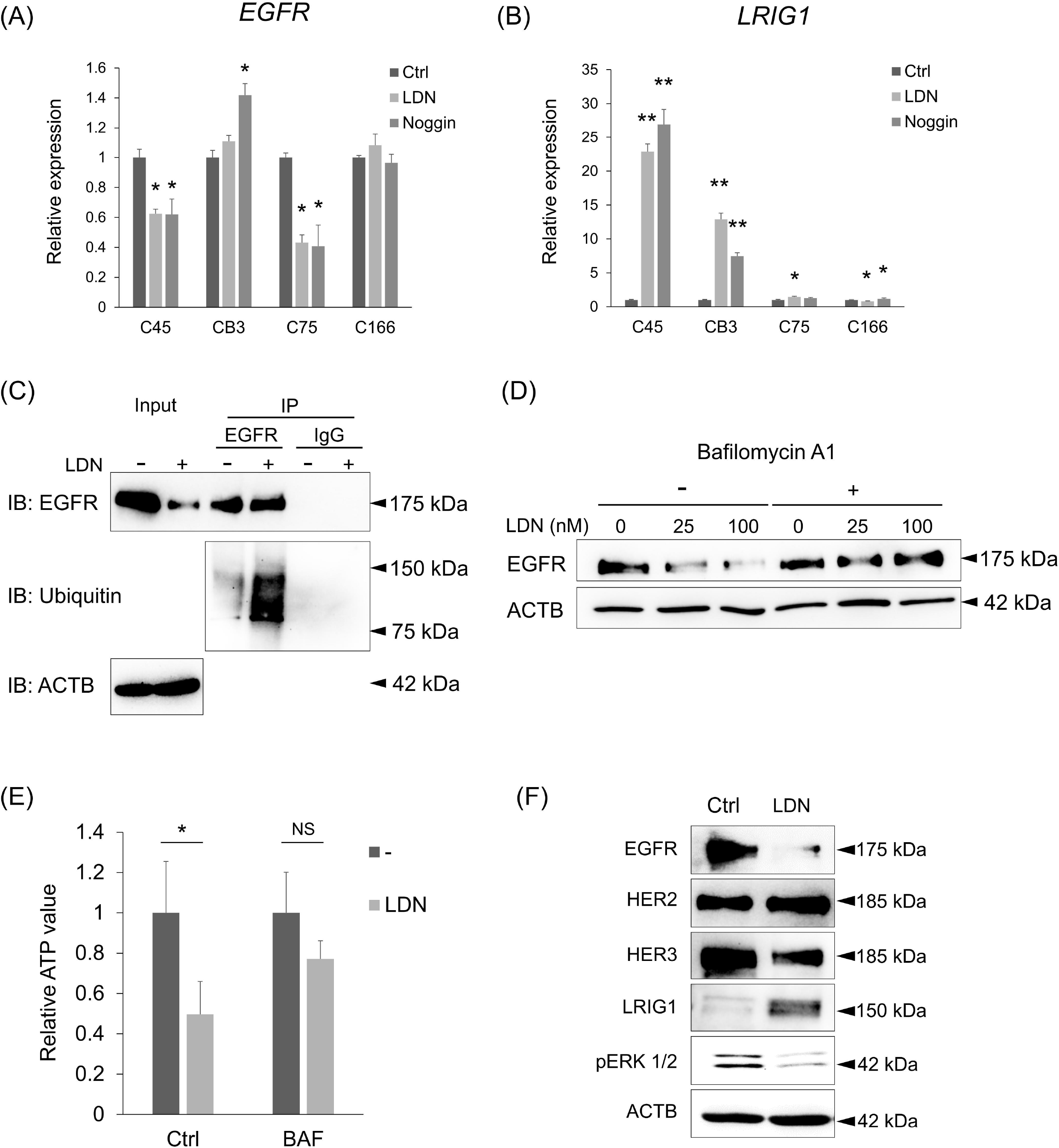
BMP inhibition induces LRIG1-mediated degradation of EGFR. **A-B**, Gene expression of *EGFR* (A) and *LRIG1* (B) in CRC organoids treated with LDN or Noggin. The expression level was normalized to control (Ctrl) in each organoid. N = 3 for each condition. Data are presented as mean + SD. Statistical comparisons were made for each control condition. * P < 0.05, ** P < 0.001, t-test with Bonferroni correction. **C**, Co-immunoprecipitation analysis of lysates from C45 organoids treated with 0.1 μM LDN (LDN) comparing to untreated control (Ctrl). Samples were immunoblotted with EGFR, ubiquitin, and ACTB antibodies. Input lysates were used as loading controls. **D**, Immunoblotting analysis of C45 organoid cultured for 48 h with LDN at the indicated concentration (nM) in the presence (+) or absence (-) of 5 nM Bafilomycin A1. Proteins were detected with indicated antibodies. **E**, Cell viability assay results for C45 organoids. Organoids were treated with 0.1 μM LDN with or without 5 nM Bafilomycin A1 (BAF). Data are presented as mean + SD. N = 3. * P < 0.05, t-test with Bonferroni correction. **F**, Immunoblotting analysis of C45 organoid cultured in the presence (LDN) or absence (Ctrl) of LDN for 48 h. Proteins were detected with indicated antibodies.

### 3.4 LRIG1 induction by BMP inhibition suppresses the growth of CRC organoids

In the 18 CRC organoid lines, *LRIG1* induction by LDN was inversely correlated with the relative growth of organoids with vs. without LDN in the medium (Fig. 4A), suggesting that *LRIG1* is involved in LDN-induced growth suppression. However, induction of the ERK negative feedback factor *DUSP5*, which is reportedly involved in the growth suppression of CRC cell lines treated with LDN ^8^, was not correlated with LDN-induced growth suppression in these 18 CRC organoids (Fig. S4). Next, induction of the Wnt target genes was evaluated because the Wnt pathway is crucial in CRC pathogenesis. In addition, *LRIG1* is a marker of intestinal and colorectal epithelial stem cells which have high Wnt activity ^24,28^, and can be regulated by Wnt via MYC ^29^. Induction of leucine-rich repeat containing G protein-coupled receptor (*LGR5*), a Wnt target stem cell marker gene, was correlated with organoid growth under LDN treatment, and the induction of *LRIG1* and *LGR5* was highly correlated (Fig. S5). However, the induction of two major Wnt target genes, *AXIN2* and *MYC*, was not correlated with the growth of CRC organoids treated with LDN (Fig. S4). These results suggest that *LRIG1* induction is independent of Wnt activation and is regulated by the same mechanism as *LGR5*.

**FIGURE 4.**
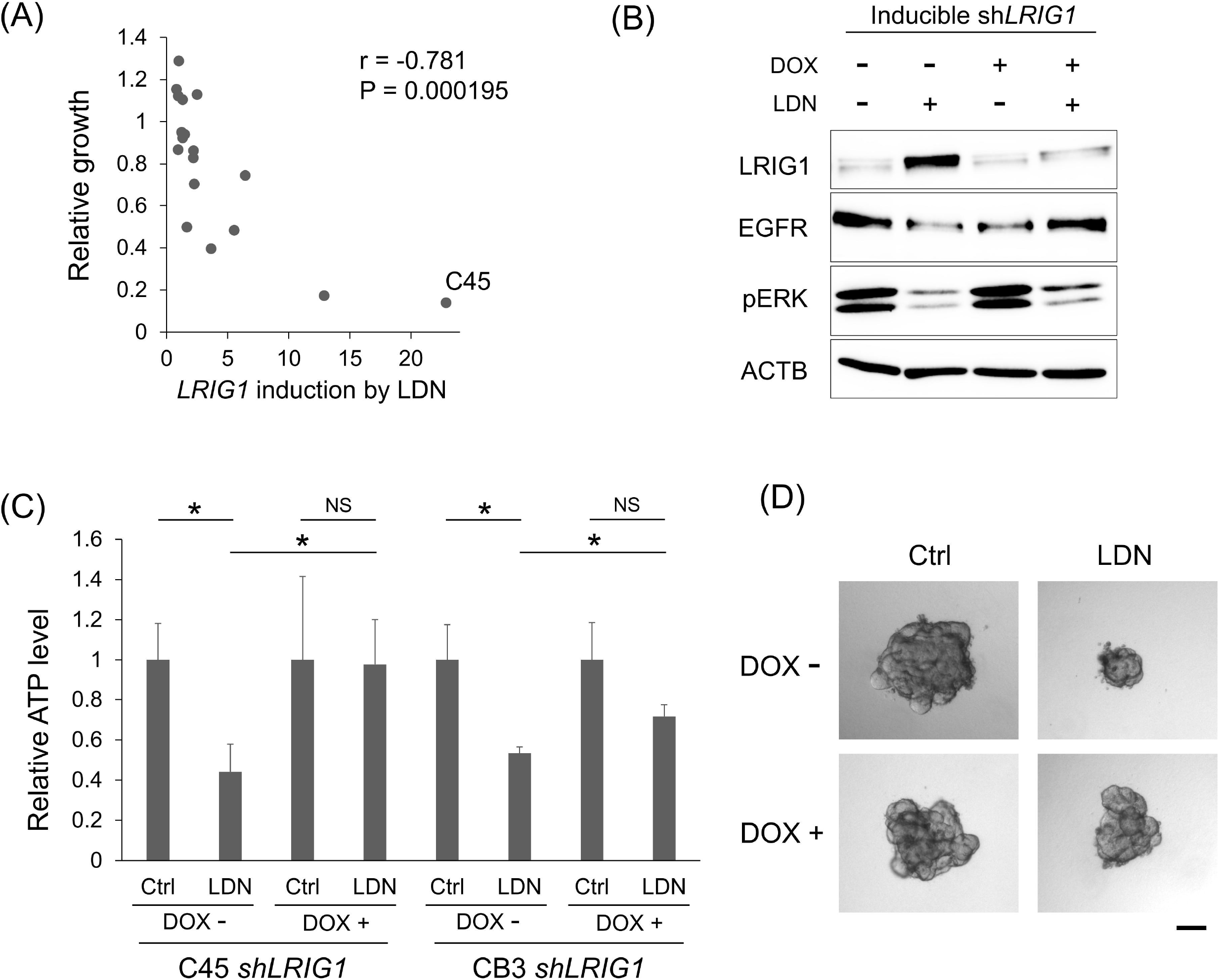
LRIG1 induction by BMP inhibition is involved in the growth suppression of CRC organoids. **A**, Scatter plot showing the correlation of the growth and LRIG1 induction in the presence of 0.1 μM LDN relative to LDN free condition. Each dot represents individual CRC organoid line (N = 18). r, Spearman’s rank correlation coefficient; P = 0.000195, Spearman’s rank correlation test. **B**, Immunoblotting analysis of C45 organoid with inducible *shLRIG1* cultured for 48 h with LDN and at the indicated concentration (nM) in the presence (+) or absence (-) of doxycycline (Dox). Proteins were detected with indicated antibodies. **C**, Cell viability assay results for C45 and CB3 organoids with inducible *shLRIG1*. Organoids were treated with 0.1 μM LDN in the presence (+) or absence (-) of doxycycline (Dox) for 5 days. Data are presented as mean + SD. N = 5. * P < 0.05, t-test with Bonferroni correction. **D**, Representative bright field images of C45 organoids with inducible *shLRIG1*. Organoids were treated with 0.1 μM LDN in the presence (+) or absence (-) of doxycycline (Dox) for 5 days. Scale bars: 100 μm.

To confirm the involvement of LRIG1 in LDN-induced growth suppression, *LRIG1* was knocked down by inducible shRNA in C45 organoids, and we found that *LRIG1* induction by LDN was the most robust (Fig. 4A). *LRIG1* knockdown (*LRIG1* KD) abrogated the downregulation of EGFR by LDN in the C45 organoids (Fig. 4B). Accordingly, the inhibition of ERK phosphorylation was slightly reversed. In addition, growth suppression by LDN was inhibited by the *LRIG1* KD in CRC organoids (Fig. 4C-D). Collectively, the growth inhibition of CRC organoids by the BMP inhibitor was, at least in part, mediated by the induction of the EGFR negative regulator, LRIG1.

### 3.5 Combined inhibition of BMP and MEK cooperatively suppresses the growth of MEK-dependent CRC organoids

As mentioned above (Fig. 2D-F), the growth–inhibitory effect of BMP inhibition was attenuated in the GF-rich microenvironment, possibly due to the strongly stimulated signaling pathways, such as MEK/ERK. The potential of CRC cells to respond to BMP inhibition is attenuated or even masked by the presence of excess growth factors. Therefore, we investigated whether BMP inhibition combined with MEK inhibition exhibited an additional effect in LDN-sensitive CRC organoids. Trametinib, a specific FDA-approved MEK inhibitor, was used as a single agent to treat the 15 CRC organoids. Trametinib treatment resulted in diverse growth inhibitory effects on these organoids under GF-free conditions (Fig. 5A). The growth inhibitory effects of LDN and trametinib were positively correlated (Fig. 5B). The organoids were then treated with trametinib and LDN. As expected, LDN-sensitive organoid lines exhibited an additive effect of these two inhibitors (Fig. 5C upper). Western blot analysis confirmed that ERK phosphorylation was suppressed more by the combination treatment than by each single agent (Fig. 5D). Of note, organoids resistant to LDN failed to show a clear combination effect (Fig. 5C lower). The combined effect was confirmed using LDN treatment with an additional MEK inhibitor, PD 0325901 (Fig. 5E). These results indicate the heterogeneity of the dependency on MEK/ERK signaling among CRC organoid lines and support the association of the LDN effect with *in vitro* MEK dependency.

**FIGURE 5.**
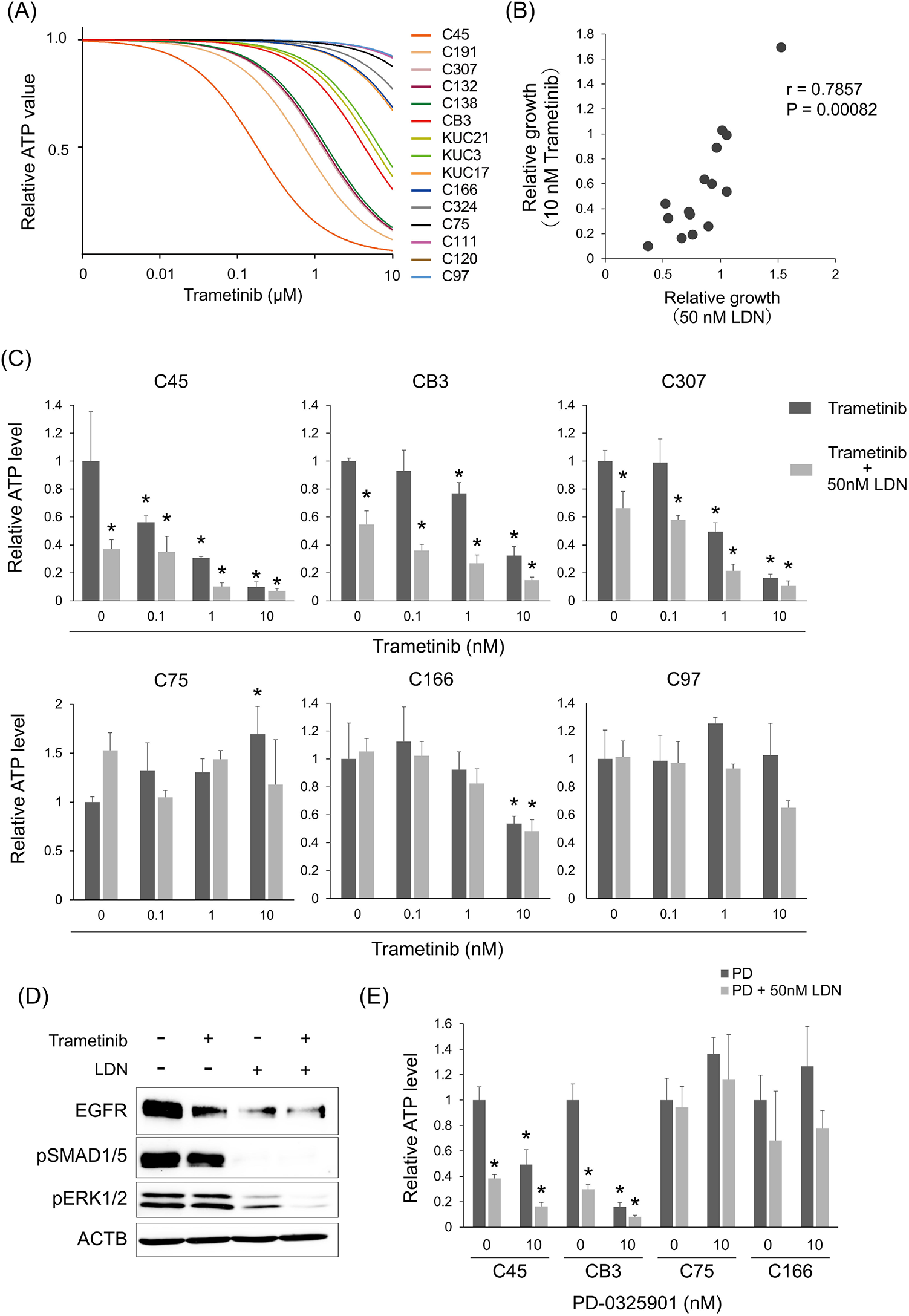
Combined inhibition of BMP and MEK cooperatively suppresses the growth of MEK-dependent CRC organoids. **A**, Dose–response curves of trametinib in 15 CRC organoid lines cultured in the GF-free medium. **B**, Scatter plot showing the correlation of the organoid growth treated with 50 nM LDN and with 10 nM trametinib relative to the drug-free condition. Each dot represents individual CRC organoid line (N = 15). r, Spearman’s rank correlation coefficient; P = 0.00082, Spearman’s rank correlation test. **C**, Cell viability assay results for MEK-dependent (upper) and -independent (lower) organoids. Organoids were treated with trametinib at indicated dose in the absence (dark gray) or presence (light gray) of LDN for 5 days. Data are presented as mean + SD. N = 3. * P < 0.05 compared with drug-free control, one-way ANOVA with post-hoc Tukey test. **D**, Immunoblotting analysis of C45 organoid cultured for 48 h with LDN and trametinib. Proteins were detected with indicated antibodies. **E**, Cell viability assay results for MEK-dependent (C45 and CB3) and -independent (C75 and C166) organoids. Organoids were treated with 10 nM PD-0325901 in the absence (dark gray) or presence (light gray) of LDN for 5 days. Data are presented as mean + SD. N = 3. * P < 0.05 compared with drug-free control in each organoid line, one-way ANOVA with post-hoc Tukey test.

### 3.6 LDN enhanced the growth inhibitory effect of trametinib in xenograft tumors of LDN-sensitive CRC organoids

To evaluate the ability of LDN to suppress the growth of CRC cells *in vivo*, mice with C45 xenograft tumors were treated with LDN. LDN had no inhibitory effect on tumor growth (Fig. 6A), which corresponds to the *in vitro* results obtained in GF-rich environments. The combined effects of LDN and trametinib were evaluated *in vivo*. Trametinib exhibited an enhanced growth inhibitory effect on C45 xenograft tumors when combined with LDN (Fig. 6B). Similarly, the growth of CB3 xenograft tumors was suppressed more efficiently by trametinib combined with LDN than by trametinib alone (Fig. 6B). However, LDN alone had no effect (Fig. 6B). Notably, the combined effect of LDN and trametinib was not observed in C166 and C75 organoids, reflecting the results of the *in vitro* organoid growth assay (Fig. 6C). Therefore, in CRC cells that are more dependent on MEK/ERK pathway activation, the simultaneous inhibition of the BMP and MEK pathways can be a potent therapeutic target.

**FIGURE 6.**
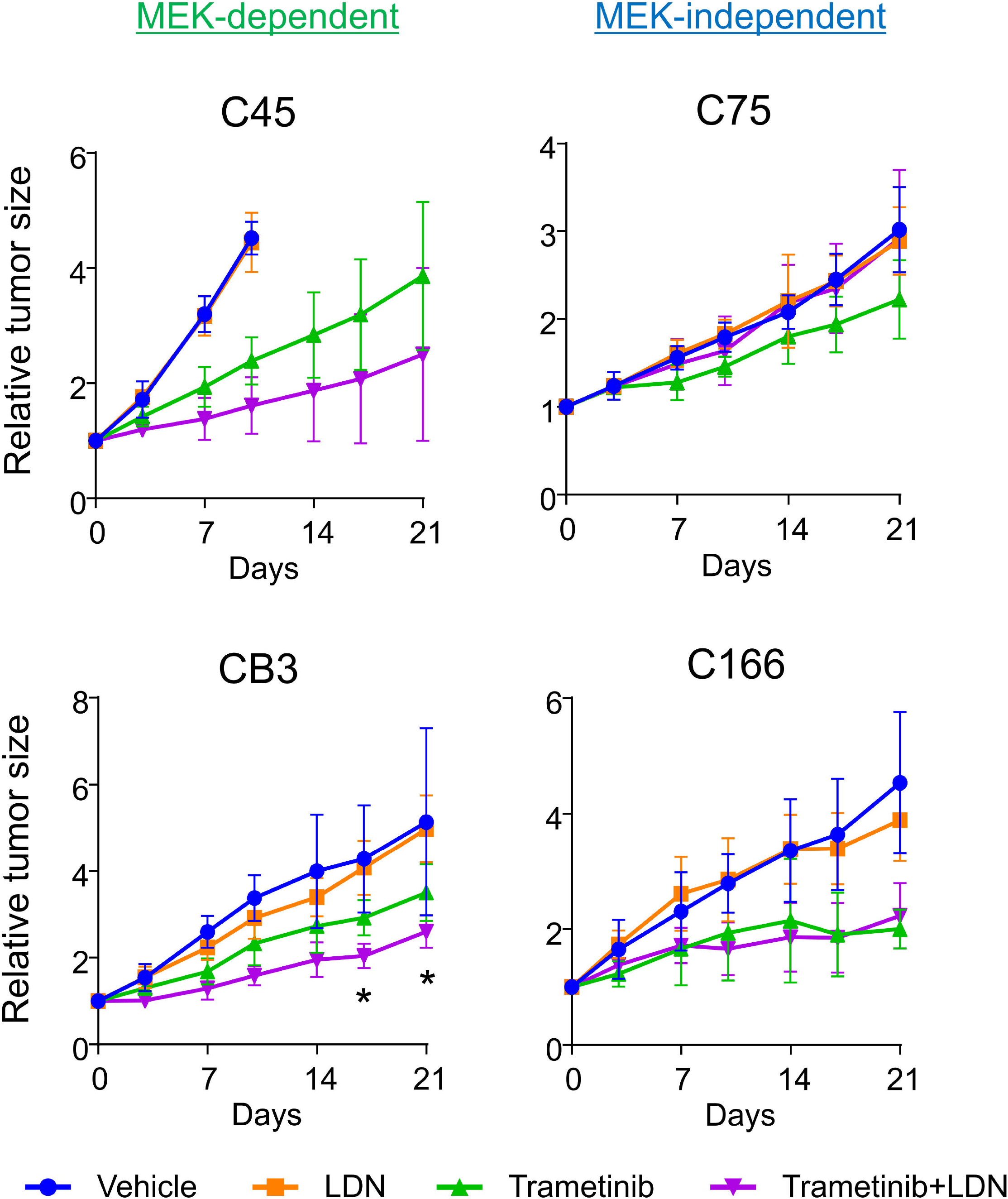
LDN enhanced the growth inhibitory effect of trametinib in xenograft tumors of LDN-sensitive CRC organoids. Growth curves of subcutaneous tumors generated by four CRC organoid lines (MEK-dependent group: C45 and CB3; MEK-independent group: C75 and C48). Blue, treated with vehicle; orange, LND (3 mg/kg) alone; green, trametinib (0.3 mg/kg) alone; purple, combination of LDN (3 mg/kg) and trametinib (0.3 mg/kg). Mean ± SD is shown. N = 4–6 in each treated group. *P < 0.05, trametinib mono therapy versus combination; one-way ANOVA with post-hoc Tukey test.

## 4. Discussion

In this study, we found that CRC is a BMP-rich environment and that BMP inhibition suppresses the growth of certain populations of CRC cells via LRIG1-mediated EGFR downregulation. The effect of BMP inhibition varied among CRC organoids, which could be masked by high levels of environmental growth factors. Accordingly, simultaneous inhibition of BMP and MEK resulted in a combined effect both *in vitro* and *in vivo*, especially in CRCs that are dependent on MEK activation.

Several studies have shown that the BMP/SMAD pathway regulates EGFR signaling; BMP2 suppresses EGFR signaling during chondrocyte maturation ^30^, and BMP signaling downregulates EGFR in astrocytes ^31^ and gastric cancer cells ^32^. All these previous reports have shown that the active BMP pathway suppresses EGFR, although the underlying mechanisms are unknown. In the present study, we found that, contrary to these previous findings, BMP inhibition downregulated the total EGFR level in some CRC organoids. We demonstrated that one possible mechanism of this phenomenon is the post-transcriptional downregulation—not transcriptional regulation—of the EGFR protein by LRIG1. In the intestine and colon, LRIG1 is a marker for epithelial stem cells ^28^ and a subpopulation of the interstitial cells of Cajal ^33^. Functionally, LRIG1 induces Cbl-dependent ubiquitination of the ErbB receptor family, which induces lysosomal degradation ^24^. Therefore, LRIG1 can serve as a tumor suppressor in many cancer types. The expression of LRIG1 is correlated with a better prognosis in several cancers, including non-small cell lung cancer ^34,35^, breast cancer ^24,36^, and hepatocellular carcinoma ^37^. Although its expression is significantly lower in CRC than in normal colorectal tissue ^24,38^, LRIG1 expression is not a prognostic biomarker in CRC at the protein and mRNA levels ^38,39^. This low LRIG1 expression might be due to the suppression of LRIG1 by BMP, which is known to suppress the expression of murine intestinal epithelial stemness genes, including *Lrig1* and *Lgr5*, via SMAD-mediated repression ^40^. This is supported by our finding that BMP inhibition upregulates the expression of *LRIG1* and *LGR5* in CRC organoids. In addition, the degree of *LRIG1* induction by BMP inhibition was correlated with sensitivity to LDN and combination therapy with LDN and trametinib *in vitro* and *in vivo*. Whether BMP is a tumor suppressor or promoter in CRC is controversial ^5–8^. Such complexities in BMP signaling have been described in various cancers ^9,10^. As CRC organoids exhibited varying responses to BMP inhibition in our study, one possible explanation for the controversy regarding the role of BMP in CRC is the heterogeneous nature of the disease. BMP is likely a tumor promoter in a certain population of CRC and can be a therapeutic target in such CRC cases.

Targeting the EGFR/RAS/RAF/MEK pathway is a major strategy for the treatment of advanced CRC. Recently, combination therapies targeting more than one molecule in the same pathway have been used in clinical trials, such as targeting EGFR and MEK in KRAS wild-type patients ^41^ and targeting EGFR and BRAF in BRAF-mutant patients ^42^. BMP inhibition may be an option for indirectly suppressing EGFR in combination with MEK inhibitors. BMP inhibition is a possible candidate for clinical combination therapy, but it is essential to select potential responders carefully because of the heterogeneous response. The mutation status of frequently mutated genes in CRC, including APC and KRAS, did not show a clear correlation with the *in vitro* response to LDN (Fig. S6). The LDN-sensitive organoid lines, C45 and CB3, were both APC and KRAS mutants. In line with our previous report that these organoid lines are partially sensitive to cetuximab ^13^, these organoids were dependent on EGFR signaling despite constitutively active *KRAS* mutations. *SMAD4* mutations, which are detected in 10% of CRC specimens according to TCGA database ^17^, inhibit SMAD mediated signaling by TGFβ and BMP ^43^. In the present study, CRC organoids with *SMAD4* mutations tended to exhibit less sensitivity to LDN and LRIG1 induction (Fig. S6), suggesting that *LRIG1* induction by LDN is a SMAD4-dependent event. Further accumulation of CRC cases is needed in the future study to evaluate the correlation between mutation status and sensitivity to LDN/trametinib combination treatment.

Taken together, these findings lead to the establishment of the BMP/SMAD pathway as a therapeutic target, especially in combination with a MEK inhibitor. Organoids prepared from patient tumor tissues are an optimal model to dissect inter-tumor heterogeneity. Notably, the organoid model allows the detection of biochemical responses to external stimuli, which cannot be assessed using single timepoint snapshot data. An “organoid assay”, such as *in vitro* LDN sensitivity or LRIG1 induction by LDN, may be useful in predicting prognosis and therapeutic efficacy. Nevertheless, a larger cohort study is necessary to clarify the heterogeneity of the CRC response to BMP.

## Supporting information

Doc S1

## Abbreviations

BMP: bone morphogenetic protein
TGF-β: transforming growth factor beta
CRC: colorectal cancer
EMT: epithelial-mesenchymal transition
EGFR: epidermal growth factor receptor
CTOS: cancer tissue-originated spheroid
LDN: LDN193189
LRIG1: leucine-rich repeats and immunoglobulin-like domains protein 1
LGR5: leucine Rich repeat containing G protein-coupled receptor 5

## Acknowledgments

The authors thank M Izutsu for secretarial assistance, A Manabe and C Saito for technical assistance. We would like to thank Editage (www.editage.com) for English language editing.

## Author Contribution

S Shimizu: Investigation, Methodology, Writing - Review & Editing, J Kondo: Conceptualization, Investigation, Validation, Visualization, Writing - Original draft preparation, Funding acquisition, K Onuma: Writing – Review & Editing, K Ota, M Kamada, Y Harada, Y Tanaka, MA Nakazawa, and Y Tamada: Formal analysis, Y Okuno: Formal analysis, Funding acquisition, RJ Coffey: Writing – Review & Editing, Y Fujiwara: Writing – Review & Editing, M Inoue: Supervision, Validation, Writing – Review & Editing, Funding acquisition.

