## Supplementary material for "Inhibition of the BMP pathway suppresses tumor growth via downregulation of EGFR in MEK/ERK-dependent colorectal cancer": Table S1

**TableS1.**

| Primer name | sequence |
| --- | --- |
| EGFR-Fw | 5’ -CCAAGGGAGTTTGTGGAGAA- 3’ |
| EGFR-Rev | 5’ -CTTCCAGACCAGGGTGTTGT- 3’ |
| LRIG1-Fw | 5’ -TGTGTCCAAGAGATGCAAGC- 3’ |
| LRIG1-Rev | 5’ -GCTTGTCCCCTGGAGTAACA- 3’ |
| ERRFI-Fw | 5’ -GACCCGATAACCATGGCCTA- 3’ |
| ERRFI-Rev | 5’ -GCACAAACCCCATTCACTGT- 3’ |
| DUSP5-Fw | 5’ -TGTCGTCCTCACCTCGCTA- 3’ |
| DUSP5-Rev | 5’ -GGGCTCTCTCACTCTCAATCTTC- 3’ |
| LGR5-Fw | 5’ -AGAATTTGCGAAGCCTTCAA- 3’ |
| LGR5-Rev | 5’ -TATTTTGTTCAGGGCCAAGG- 3’ |
| AXIN2-Fw | 5’ -AGTGTGAGGTCCACGGAAAC- 3’ |
| AXIN2-Rev | 5’ -CTTCACACTGCGATGCATTT- 3’ |
| MYC-Fw | 5’ - CAGATCAGCAACAACCGAAA- 3’ |
| MYC-Rev | 5’ - GGCCTTTTCATTGTTTTCCA- 3’ |
| ACTB-Fw | 5’ - GGACTTCGAGCAAGAGATGG- 3’ |
| ACTB-Rev | 5’ - AGCACTGTGTTGGCGTACAG- 3’ |
