## Supplementary material for "Inhibition of the BMP pathway suppresses tumor growth via downregulation of EGFR in MEK/ERK-dependent colorectal cancer": Fig. S

Figure 1

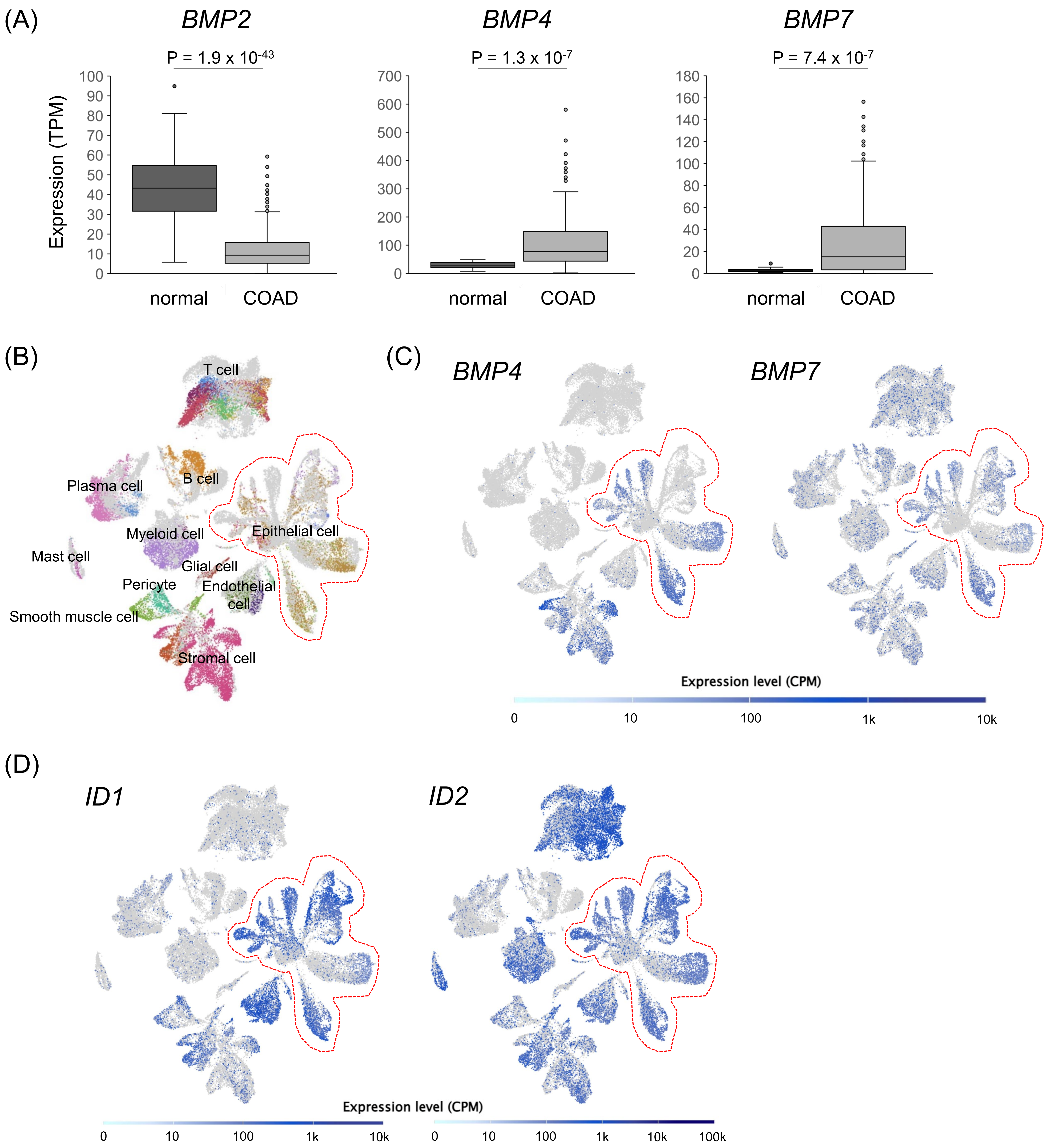

(A)
BMP2
P = 1.9 x 10-43
BMP4
P = 1.3 x 10-7
BMP7
P = 7.4 x 10-7
Expression (TPM)
normal
COAD
(B)
T cell
Plasma cell
B cell
Myeloid cell
Epithelial cell
Mast cell
Glial cell
Pericyte
Endothelial cell
Smooth muscle cell
Stromal cell
(C)
BMP4
BMP7
Expression level (CPM)
0
10
100
1k
10k
(D)
ID1
ID2
Expression level (CPM)
100k

Figure 2

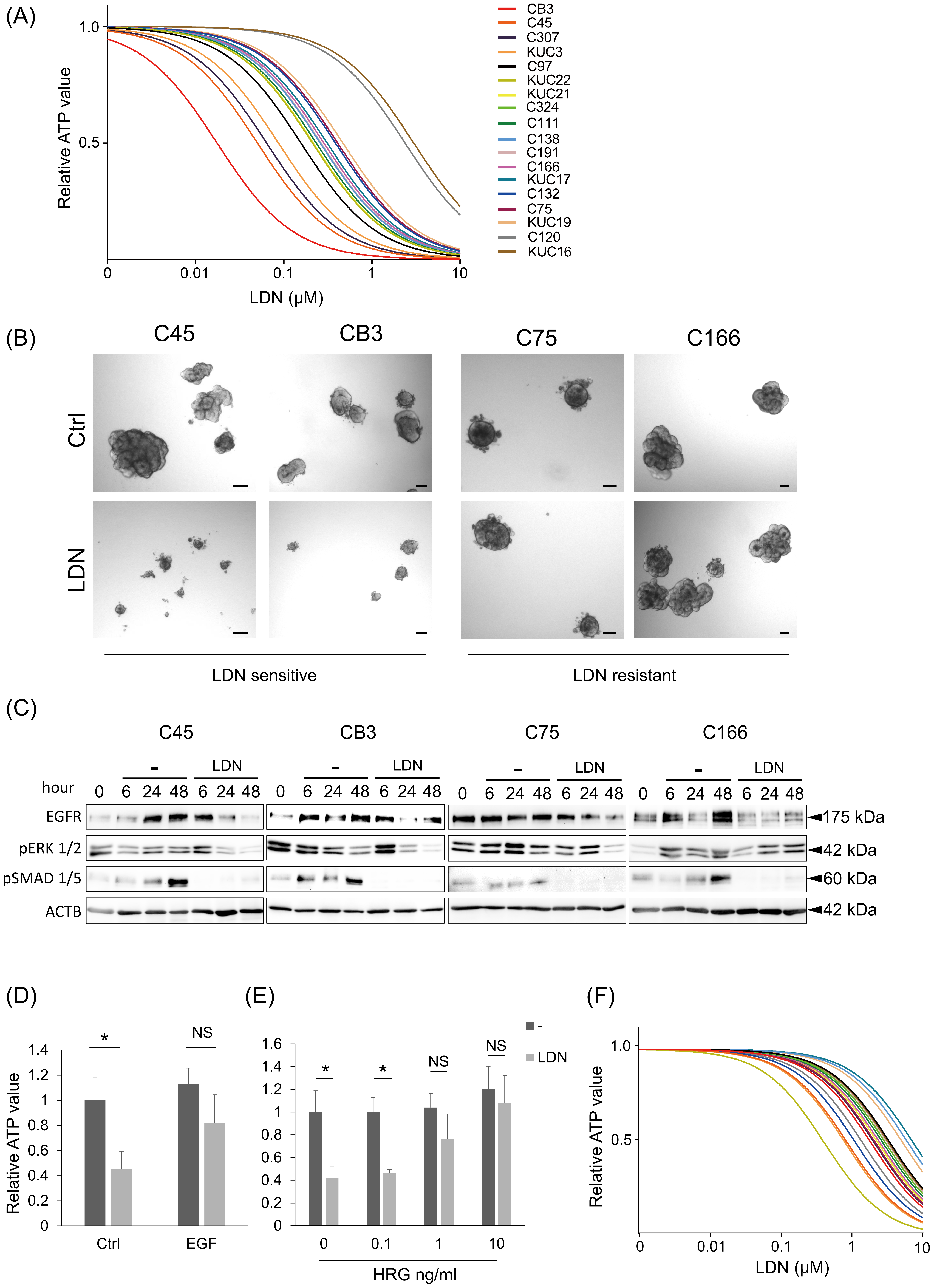

(A)
Relative ATP value
1.0
0.5
0
0.01
0.1
1
10
LDN (μM)
CB3
C45
C307
KUC3
C97
KUC22
KUC21
C324
C111
C138
C191
C166
KUC17
C132
C75
KUC19
C120
KUC16
(B)
C45
CB3
C75
C166
Ctrl
LDN
LDN sensitive
LDN resistant
(C)
C45
CB3
C75
C166
-
LDN
hour
0 6 24 48 6 24 48
EGFR
pERK 1/2
pSMAD 1/5
ACTB
◄175 kDa
◄42 kDa
◄60 kDa
◄42 kDa
(D)
Relative ATP value
*
NS
Ctrl
EGF
(E)
*
*
NS
NS
0
0.1
1
10
HRG ng/ml
-
LDN
(F)
Relative ATP value
1.0
0.5
LDN (μM)

Figure 3

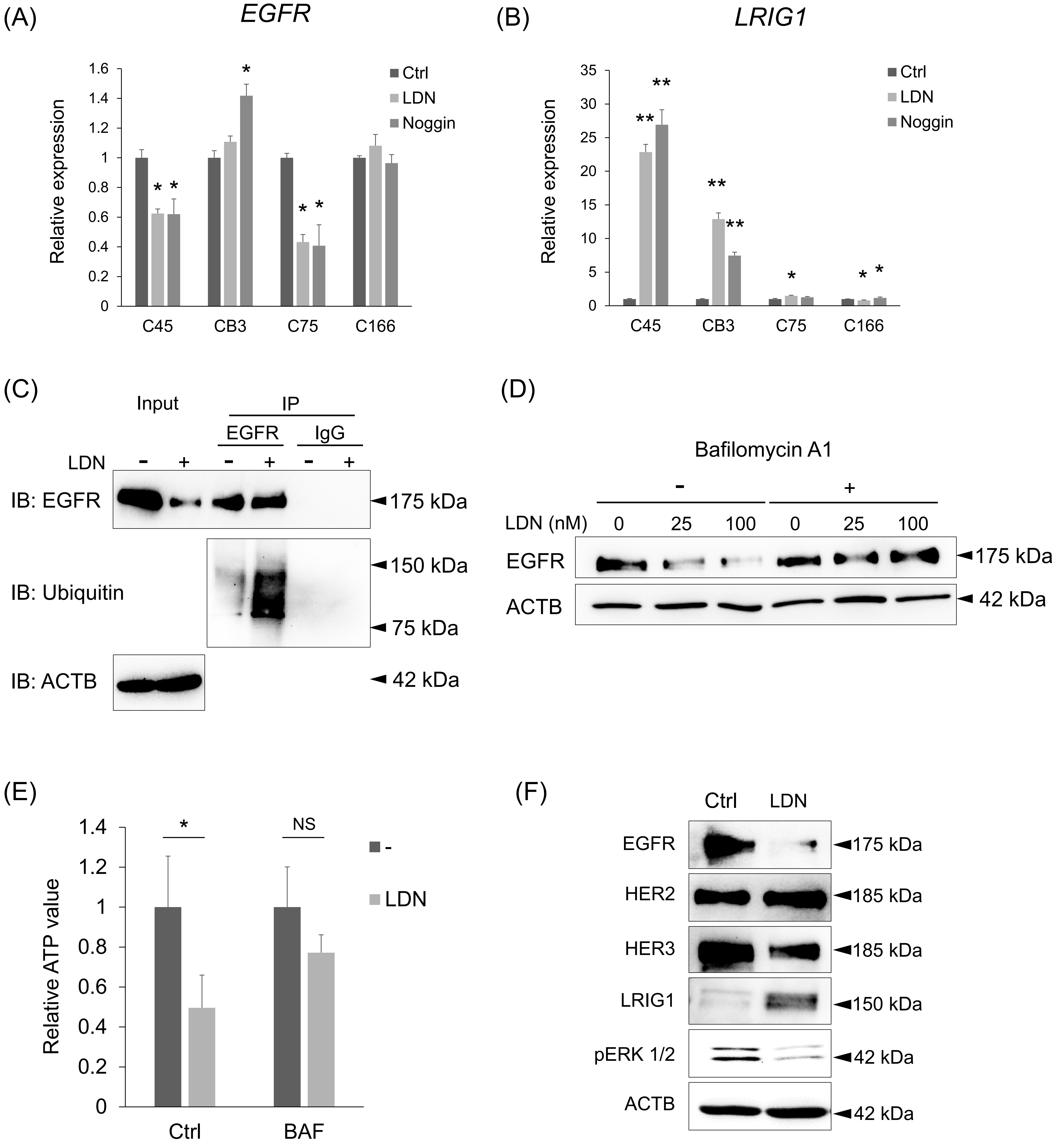
(A)
EGFR
Relative expression
Ctrl
LDN
Noggin
C45
CB3
C75
C166
(B)
LRIG1
(C)
Input
IP
EGFR
IgG
LDN
IB: EGFR
IB: Ubiquitin
IB: ACTB
175 kDa
150 kDa
75 kDa
42 kDa
(D)
Bafilomycin A1
LDN (nM)
0
25
100
EGFR
ACTB
(E)
Relative ATP value
NS
Ctrl
BAF
(F)
HER2
HER3
LRIG1
pERK 1/2
185 kDa

Figure 4

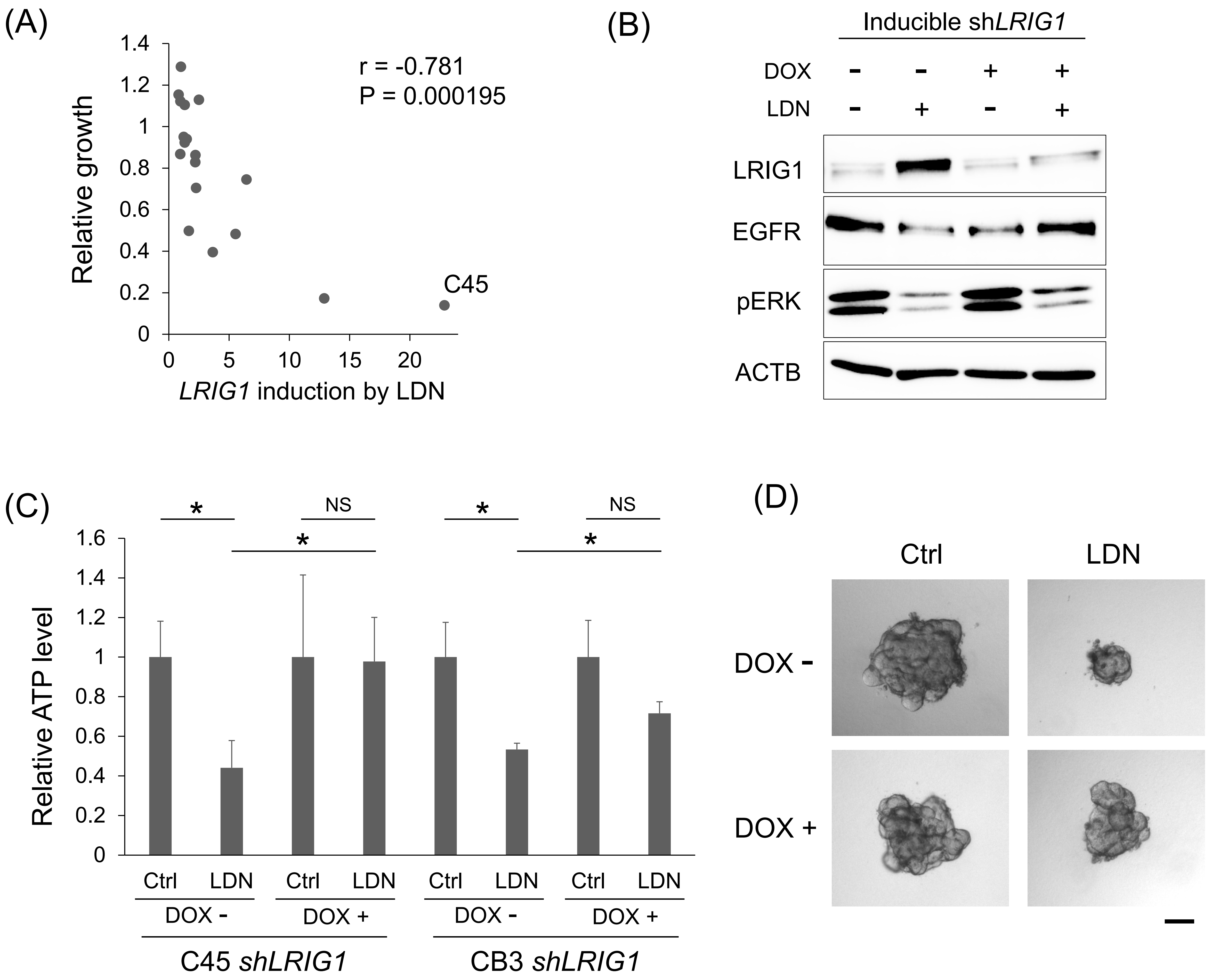
(A)
r = -0.781
P = 0.000195
Relative growth
LRIG1 induction by LDN
C45
(B)
Inducible shLRIG1
DOX
LDN
LRIG1
EGFR
pERK
ACTB
(C)
Relative ATP level
NS
Ctrl
LDN
DOX -
DOX +
C45 shLRIG1
CB3 shLRIG1
(D)
Ctrl
LDN
DOX -
DOX +

Figure 5

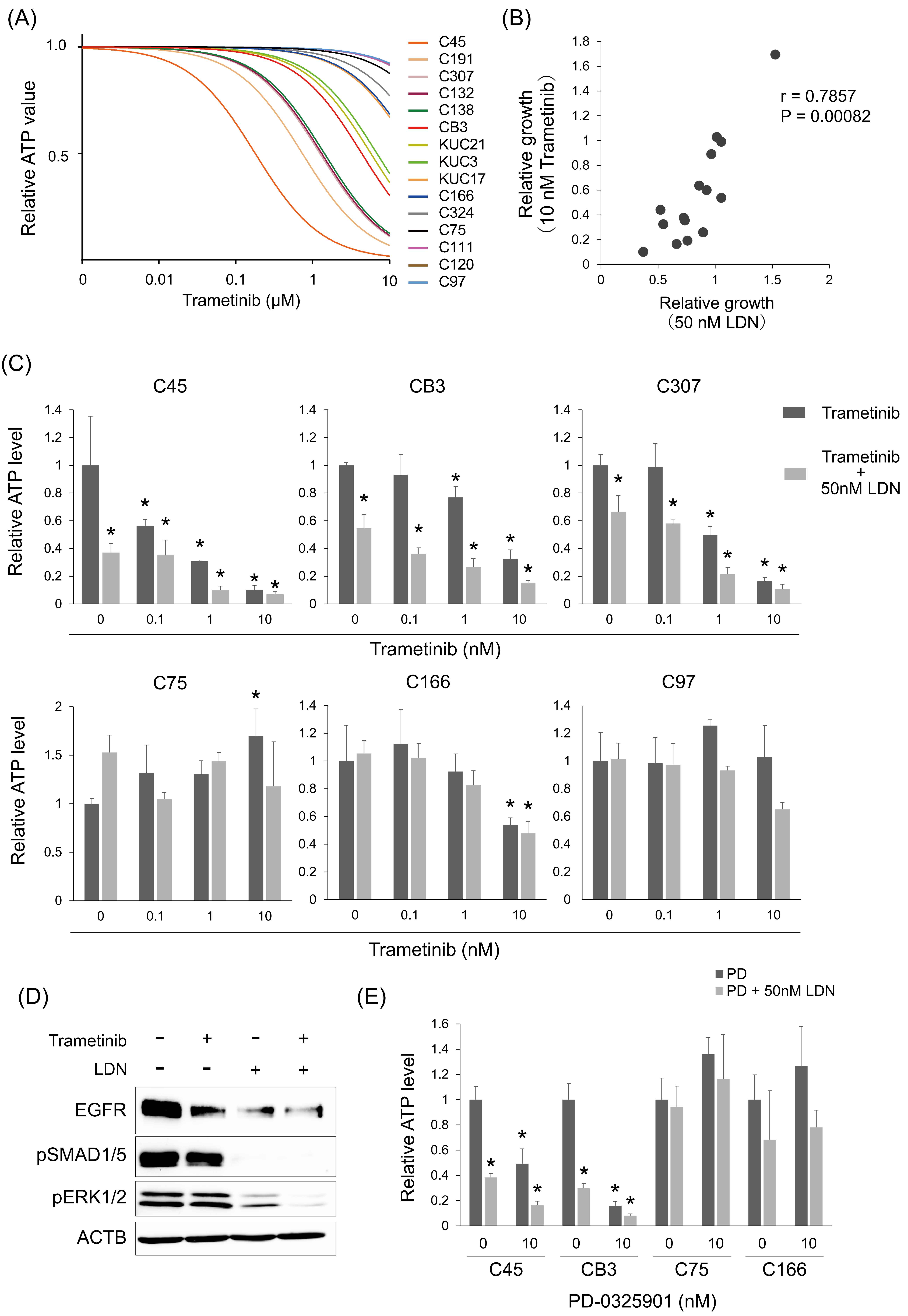

(A)
1.0
0.5
Relative ATP value
0
0.01
0.1
1
10
Trametinib (μM)
C45
C191
C307
C132
C138
CB3
KUC21
KUC3
KUC17
C166
C324
C75
C111
C120
C97
(B)
Relative growth (10 nM Trametinib)
r = 0.7857
P = 0.00082
Relative growth (50 nM LDN)
(C)
C45
CB3
C307
Trametinib
Trametinib + 50nM LDN
Relative ATP level
Trametinib (nM)
C75
C166
C97
Relative ATP level
Trametinib (nM)
(D)
Trametinib
LDN
EGFR
pSMAD1/5
pERK1/2
ACTB
(E)
PD
PD + 50nM LDN
Relative ATP level
C45
CB3
C75
C166
PD-0325901 (nM)

Figure 6

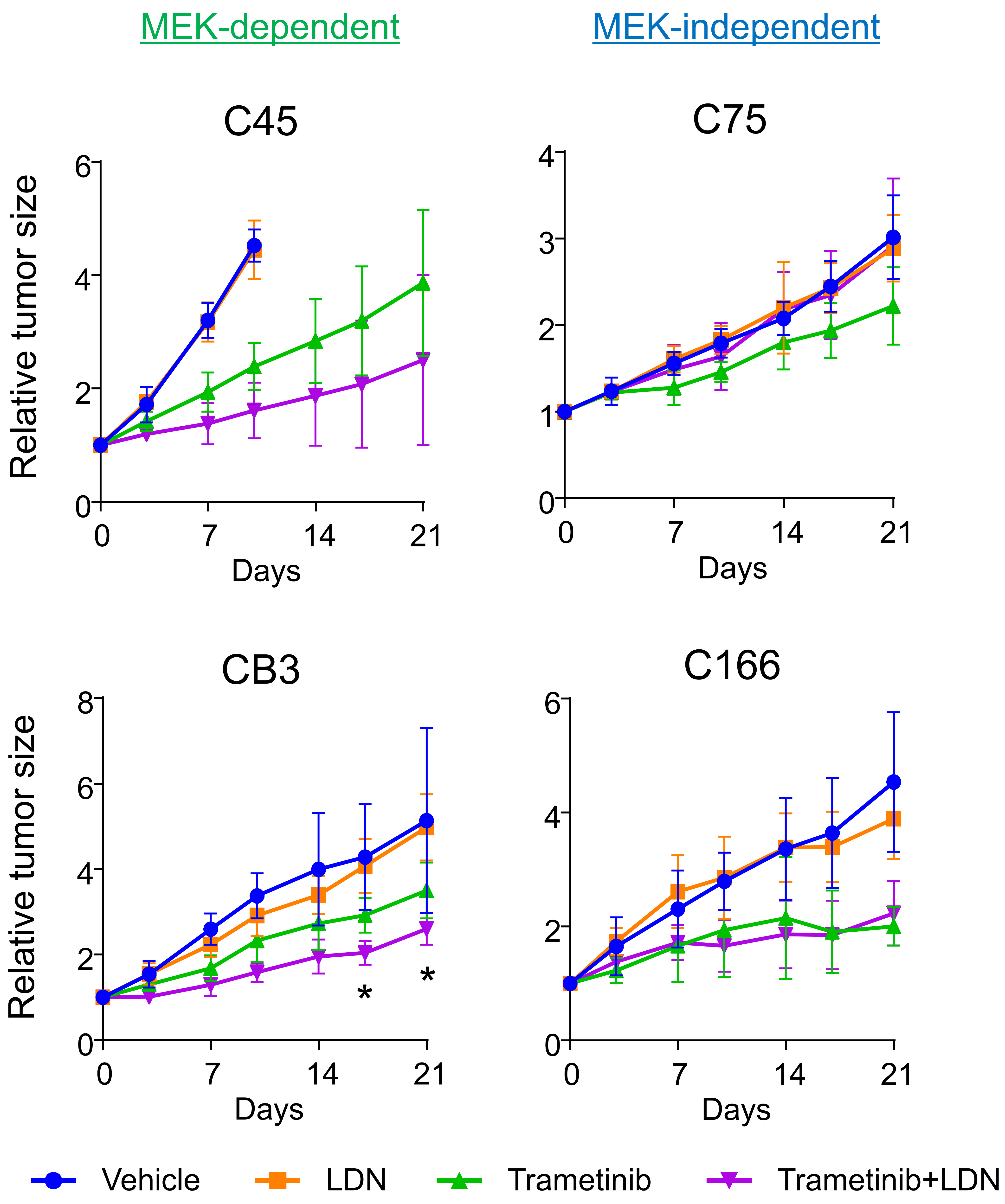

MEK-dependent
MEK-independent
C45
C75
CB3
C166
Relative tumor size
Days
0
7
14
21
*
*
Vehicle
LDN
Trametinib
Trametinib+LDN

Figure S1

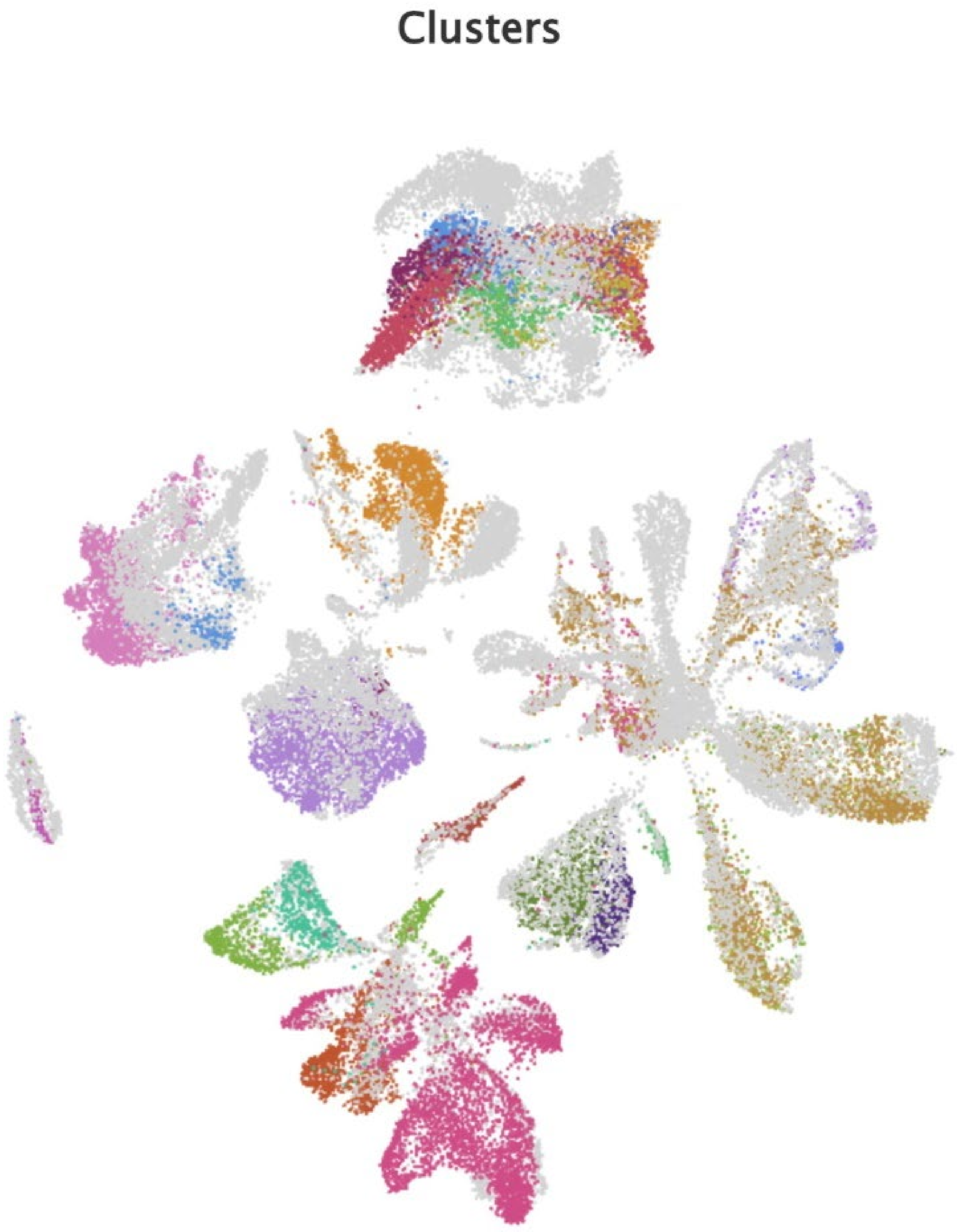


FIGURE S1. Uniform Manifold Approximation and Projection (UMAP) for the single-cell RNA sequencing data of 9 CRC patients' specimens. Detailed cell types corresponding to each cluster were designated on the UMAP (related to Fig. 1B).

Figure S2

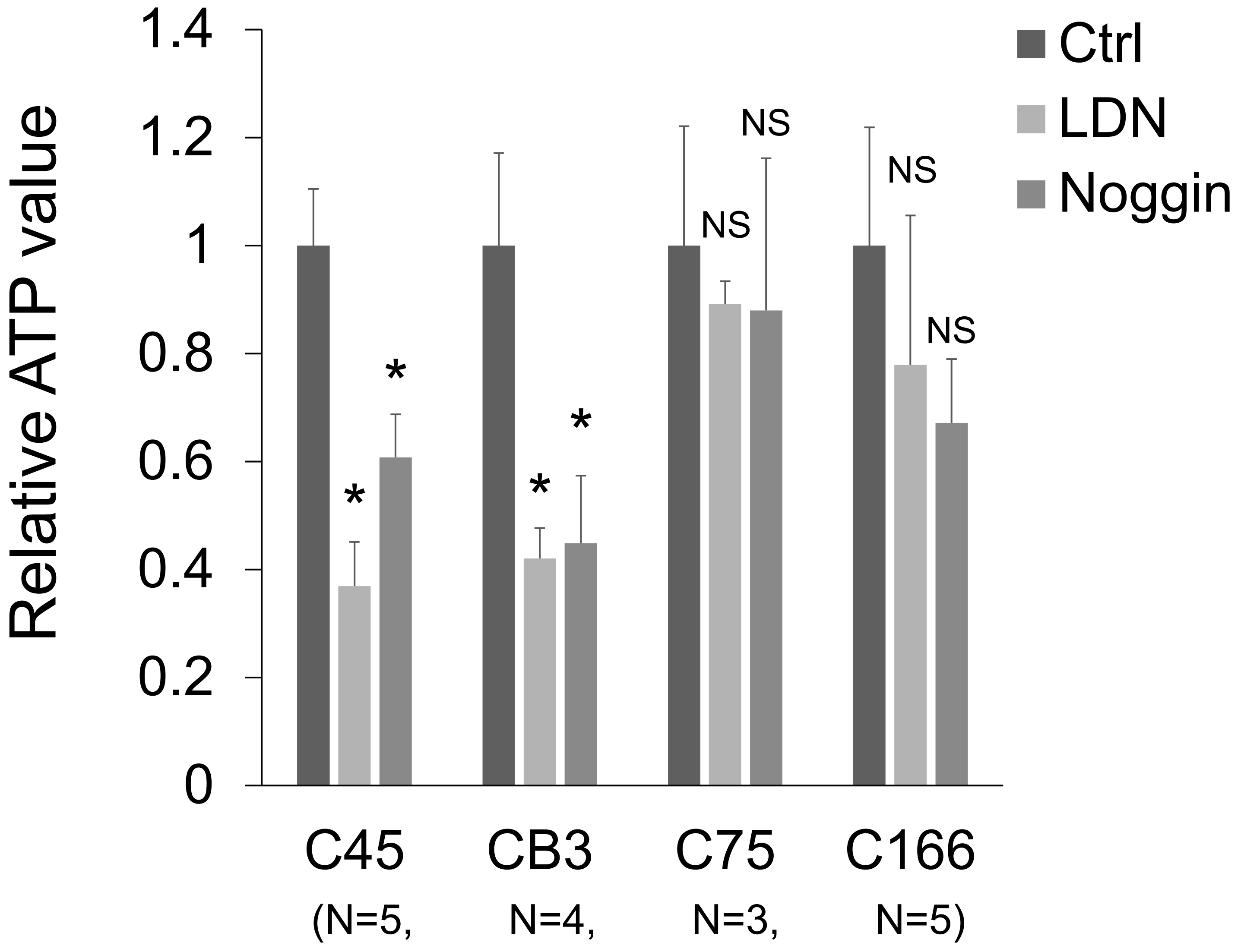


FIGURE S2. Cell viability assay results for CRC organoids. Organoids were treated with 0.1 μM LDN or 100 ng/ml. The number of replicates is indicated below the organoid names. Data are presented as mean + SD. Statistical comparisons were made for each control condition. * P < 0.05; NS, not significant, t-test with Bonferroni correction.

Figure S3

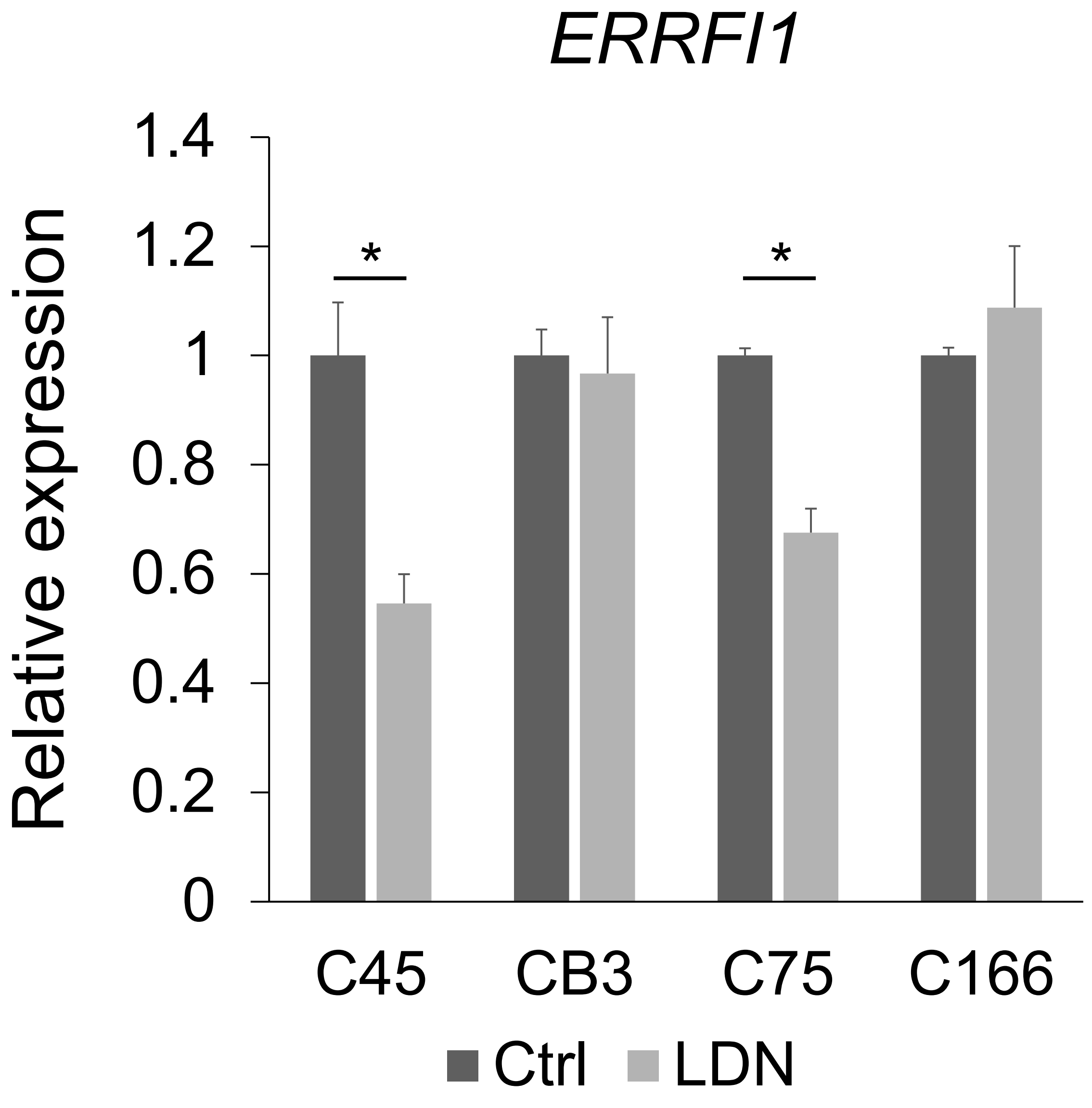


FIGURE S3. Gene expression of *ERRFI1* in CRC organoids treated with LDN. The expression level was normalized to control (Ctrl) in each organoid. N = 3 for each condition. Data are presented as mean + SD. Statistical comparisons were made for each control condition. * P < 0.001, t-test with Bonferroni correction.

Figure S4

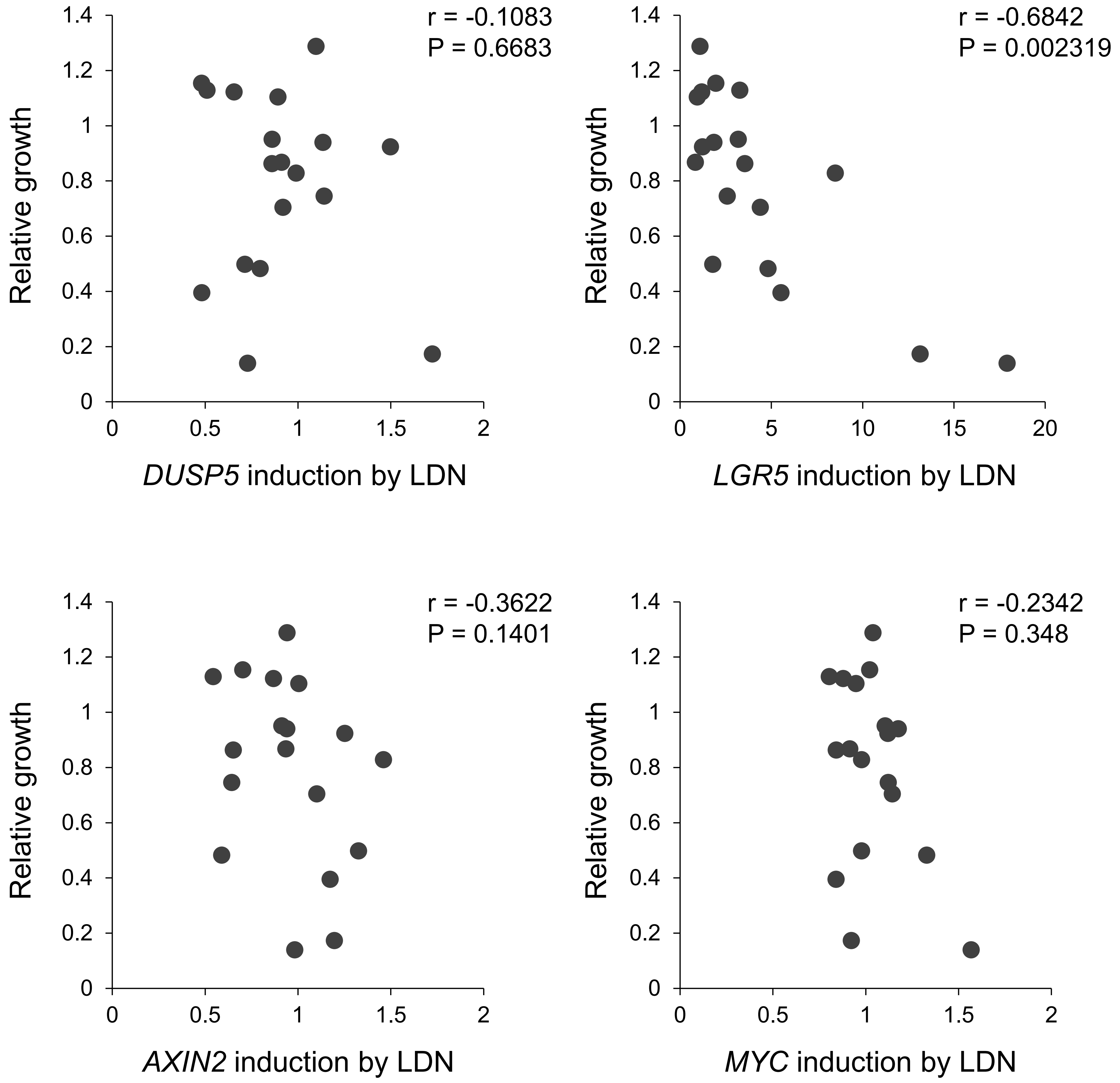


FIGURE S4. Scatter plot showing the correlation of organoid growth and *DUSP5, LGR5, AXIN2, and MYC* induction in the presence of 0.1 μM LDN relative to the LDN-free condition. Each dot represents an individual CRC organoid line (N=18). r, Spearman’s rank correlation coefficient; P value, Spearman’s rank correlation test.

Figure S5

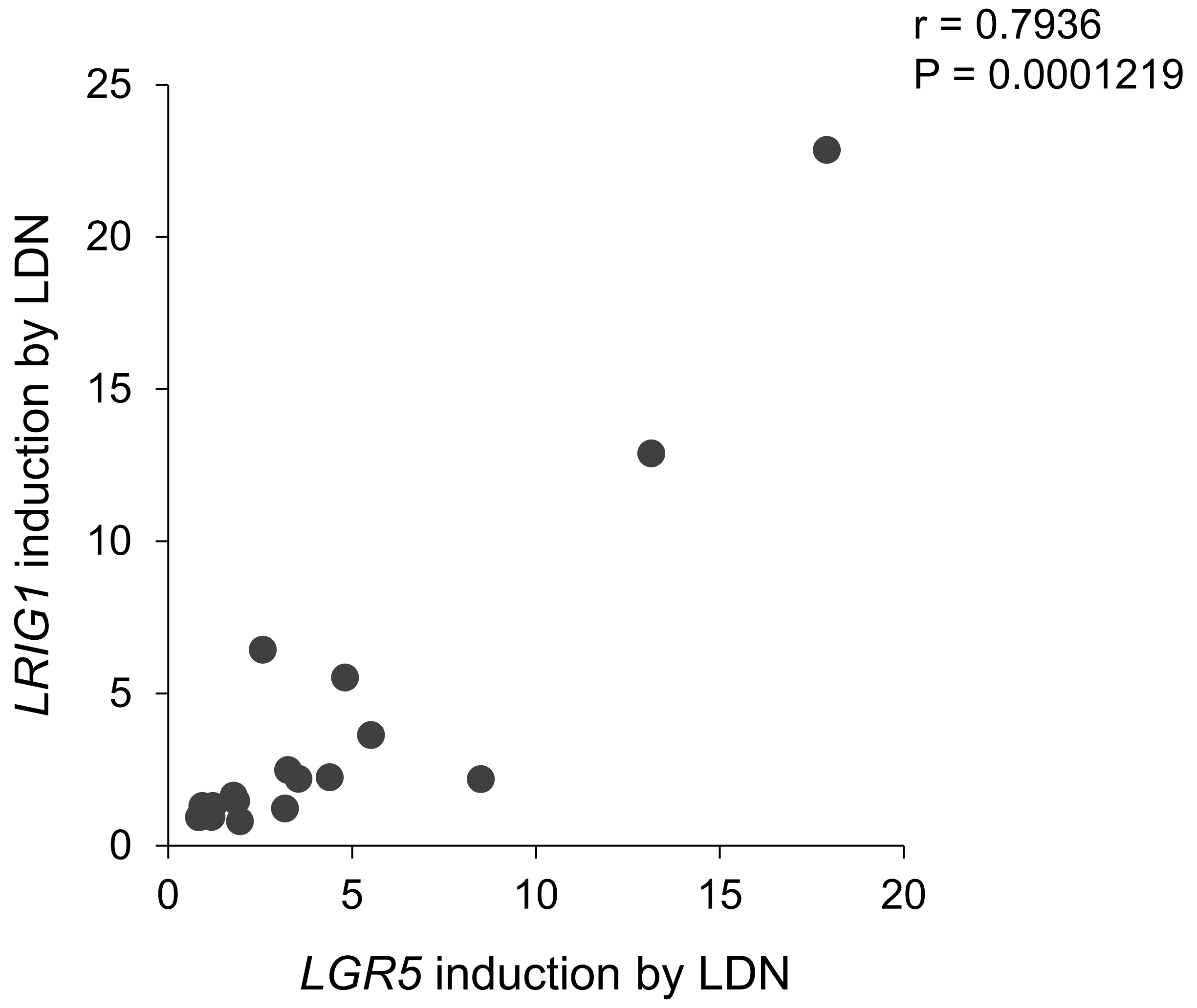


FIGURE S5. Scatter plot showing the correlation of *LRIG1 and LGR5* induction in the presence of 0.1 μM LDN relative to the LDN-free condition. Each dot represents an individual CRC organoid line (N=18). r, Spearman's rank correlation coefficient; P value, Spearman's rank correlation test.

### Figure S6

| Sample ID | LDN (0.1μM) growth ratio | LDN (0.1μM) *LRIG1* induction | Mutational status of organoids: APC | TP53 | KRAS | BRAF | PIK3CA | AKT1 | SMAD4 | CTNNB1 | LTN1 | MLH1 | FBXW7 |
|---|---|---|---|---|---|---|---|---|---|---|---|---|---|
| KUC19 | 1.287525597 | 0.999455014 | Mut | WT | Mut | WT | WT | WT | Mut | WT | WT | WT | WT |
| C75 | 1.153373842 | 0.804953166 | Mut | WT | Mut | WT | WT | Mut | WT | WT | WT | WT | WT |
| C132 | 1.128608715 | 2.487417965 | Mut | WT | WT | WT | Mut | WT | WT | WT | Mut | WT | WT |
| KUC16 | 1.122026864 | 0.951780056 | Mut | Mut | WT | WT | WT | WT | Mut | WT | WT | WT | WT |
| C120 | 1.103893588 | 1.311911342 | WT | WT | WT | Mut | Mut | WT | WT | WT | WT | WT | WT |
| KUC17 | 0.950358478 | 1.228127066 | Mut | Mut | Mut | WT | WT | WT | Mut | WT | Mut | WT | WT |
| C166 | 0.939583878 | 1.473004254 | Mut | Mut | WT | WT | WT | WT | WT | WT | WT | Mut | Mut |
| C191 | 0.923064563 | 1.312232253 | WT | Mut | WT | WT | WT | WT | Mut | Mut | WT | WT | WT |
| C138 | 0.867100823 | 0.939548321 | Mut | Mut | Mut | WT | WT | WT | Mut | WT | WT | WT | WT |
| C111 | 0.862624559 | 2.19993762 | Mut | Mut | WT | WT | WT | WT | WT | WT | WT | WT | WT |
| KUC21 | 0.828055279 | 2.189574955 | Mut | Mut | Mut | WT | WT | WT | WT | WT | WT | WT | WT |
| KUC22 | 0.745287972 | 6.438410824 | Mut | Mut | WT | WT | Mut | WT | WT | WT | WT | WT | WT |
| C97 | 0.704131701 | 2.256117353 | WT | Mut | Mut | WT | Mut | WT | WT | WT | WT | WT | WT |
| KUC3 | 0.497852501 | 1.646142092 | Mut | Mut | WT | WT | WT | WT | WT | WT | WT | WT | WT |
| C324 | 0.482312897 | 5.523302072 | Mut | WT | WT | WT | WT | WT | WT | WT | WT | WT | WT |
| C307 | 0.394814617 | 3.634832505 | Mut | Mut | Mut | WT | Mut | WT | WT | WT | WT | WT | WT |
| CB3 | 0.173005112 | 12.88706856 | Mut | Mut | Mut | WT | WT | WT | WT | WT | WT | WT | WT |
| C45 | 0.139401236 | 22.85927219 | Mut | Mut | Mut | WT | WT | WT | WT | WT | WT | WT | WT |

FIGURE S6. Growth ratio and *LRIG1* expression under treatment with 0.1 μM LDN relative to vehicle control for each of 18 CRC organoid lines are presented along with the status of detected mutation. The list is sorted by growth ratio.
