## Supplementary material for "Inhibition of the BMP pathway suppresses tumor growth via downregulation of EGFR in MEK/ERK-dependent colorectal cancer": Doc S1

**Supplementary Information**

**Supplementary Materials and Methods**

**Western blotting and co-immunoprecipitation**

Approximately 2,000 organoids (φ40–100 μm) were cultured in a 24-well plate with 2 mL GF-free medium containing 2.5% Matrigel GFR, and after 6 h of starvation, treated with the indicated dose of LDN, trametinib, or bafilomycin A1, and collected at the indicated time points. Western blotting was performed as previously described ^1^. Primary antibodies against EGFR (clone D38B1), pEGFR (Tyr1068) (clone D7A5), pERK (pp44/42 MAPK, Thr202/Tyr204) (clone D13.14.4E), and pSMAD1/5 (Ser463/465) (41D10) were obtained from Cell Signaling Technology (Danvers, MA, USA) and anti-ACTB (clone AC-15) from Sigma-Aldrich.

For immunoprecipitation, organoids, either pre-treated with LDN for 48 h or non-treated, were exposed to EGF 20 ng/mL for 30 min before collection. Organoids were then homogenized and lysed with TNE-T buffer (10 mM Tris-HCl pH 7.4, 1 mM EDTA, 150 mM NaCl, 1% Triton) containing Roche cOmplete™ EDTA-free protease inhibitor (Sigma-Aldrich) at 4 °C. After centrifugation at 14,000 rpm for 15 min at 4 °C, the supernatant was collected and pre-cleared with Protein G Sepharose 4 Fast Flow (Cytiva, Marlborough, MA, USA) for 1 h at 4 °C. The lysate was incubated with antibodies at 4 °C overnight, and then the antibodies were collected by incubation with Protein G Sepharose. After washing with TNE-T buffer, immunoprecipitated proteins were eluted by heating in 2x SDS-PAGE sample buffer.

**Semi-quantitative real-time PCR**

Organoids were treated with LDN or Noggin for 48 h and collected using the same method described above for western blotting. RNA was extracted using an RNeasy Mini Kit (Qiagen, Hilden, Germany). Reverse transcription was performed using Super Script III reverse transcriptase (Thermo Fisher Scientific), according to the manufacturer’s instructions. Semi-quantitative real-time PCR was performed using the Fast SYBR Green Master Mix (Thermo Fisher Scientific) with the StepOneReal-Time PCR System (Applied Biosystems, Foster City, CA, USA). The primer sequences are reported in Table S1. Gene expression was normalized to the *ACTB* signal to calculate relative expression levels using the 2ΔΔCq method ^2^. All data are expressed as the mean ± SD of three replicates.

**Database analysis**

Comparison of BMPs 2, 4, and 7 expression in CRC and normal tissues was performed using OncoDB ^3^. Gene expression in each cellular component of CRC tumors was analyzed with publicly available databases ^4^ using the Single Cell Expression Atlas website ^5^.

**Analysis of mutation status**

The mutation status of C45, C132, C138, C166, C307, C324, CB3, KUC3, KUC17, and KUC21 was evaluated by exome sequencing. For exome sequencing, DNA library preparation, capture, and sequencing were conducted by RIKEN Genesis CO., LTD., (Kanagawa, Japan). The captured libraries were prepared from the Agilent SureSelect Human All Exon v6 enrichment kit (Agilent Technologies, Santa Clara, USA) and Agilent SureSelect Target Enrichment System (Agilent Technologies, Santa Clara, USA). The enriched DNA libraries were sequenced using 151-Bp paired-end reads on the Illumina NovaSeq 6000 (Illumina, California, USA). After obtaining the raw sequencing data, adapter removal and quality control were conducted with Trimmomatic (v0.39) ^6^. Filtered reads were mapped to the hg38 reference sequence using BWA (v0.7.17) ^7^. For the mapping results, duplicated reads were flagged with Picard (v2.25.7) (http://broadinstitute.github.io/picard/) and preprocessing for tumor-only variant calling was performed with GATK (v4.2.0) ^8^. Sequence variants were called using Mutect2 ^9^. All the variants found in ToMMo 14KJPN (https://jmorp.megabank.tohoku.ac.jp/ijgvd/) with frequency > 1.0% were removed, and the final candidate somatic variants were annotated with SnpEff (5.0e) ^10^. For the rest of the organoids, mutation status has been reported in the previous reports ^11,12^.
